# Using a function-first ‘scout fragment’-based approach to develop allosteric covalent inhibitors of conformationally dynamic helicase mechanoenzymes

**DOI:** 10.1101/2023.09.25.559391

**Authors:** Jared R. Ramsey, Patrick M.M Shelton, Tyler K. Heiss, Paul Dominic B. Olinares, Lauren E. Vostal, Heather Soileau, Michael Grasso, Sara Warrington, Stephanie Adaniya, Michael Miller, Shan Sun, David J. Huggins, Robert W. Myers, Brian T. Chait, Ekaterina V. Vinogradova, Tarun M. Kapoor

## Abstract

Helicases, classified into six superfamilies, are mechanoenzymes that utilize energy derived from ATP hydrolysis to remodel DNA and RNA substrates. These enzymes have key roles in diverse cellular processes, such as genome replication and maintenance, ribosome assembly and translation. Helicases with essential functions only in certain cancer cells have been identified and helicases expressed by certain viruses are required for their pathogenicity. As a result, helicases are important targets for chemical probes and therapeutics. However, it has been very challenging to develop selective chemical inhibitors for helicases, enzymes with highly dynamic conformations. We envisioned that electrophilic ‘scout fragments’, which have been used for chemical proteomic based profiling, could be leveraged to develop covalent inhibitors of helicases. We adopted a function-first approach, combining enzymatic assays with enantiomeric probe pairs and mass spectrometry, to develop a covalent inhibitor that selectively targets an allosteric site in SARS-CoV-2 nsp13, a superfamily-1 helicase. Further, we demonstrate that scout fragments inhibit the activity of two human superfamily-2 helicases, BLM and WRN, involved in genome maintenance. Together, our findings suggest a covalent inhibitor discovery approach to target helicases and potentially other conformationally dynamic mechanoenzymes.

## Introduction

Developing chemical inhibitors for helicases, which are important targets for anti-viral and anti-cancer drugs, has been notoriously difficult^1,2^. There are at least two reasons why targeting helicases has been challenging. First, these enzymes undergo substantial conformational changes during the ATP hydrolysis cycle, a property that poses substantial difficulties for structure-guided inhibitor design^2^. Of the six helicase superfamilies (SFs), the mechanochemical cycle of SF1 and SF2 helicases are best understood^3^. For both these helicase superfamilies, the two core RecA-like domains transition between ‘open’ and ‘closed’ conformations during the ATP hydrolysis cycle^2^. For example, structural studies have revealed a ∼15 Å difference in the spacing of the RecA-like domains between the ATP analog-bound and apo states of the SF2 Hepatitis C NS3 helicase^4^. Second, although high-throughput activity-based screens have yielded hits for helicases, the vast majority of these compounds were subsequently found to be non-selective, false positives or indirect inhibitors (e.g. DNA intercalators)^1^.

We envisioned that a covalent inhibitor discovery approach for helicases could address both challenges, as these compounds would remain bound to the targets throughout the conformational changes linked to the enzymatic cycle (Figure 1A), and direct target engagement could be readily assessed using mass spectrometry (MS) techniques. The use of covalent probes to discover and target allosteric sites has been shown to be an effective strategy for other protein superfamilies that have been difficult to chemically inhibit^5,6^, such as K-Ras^7^.

**Figure 1.**
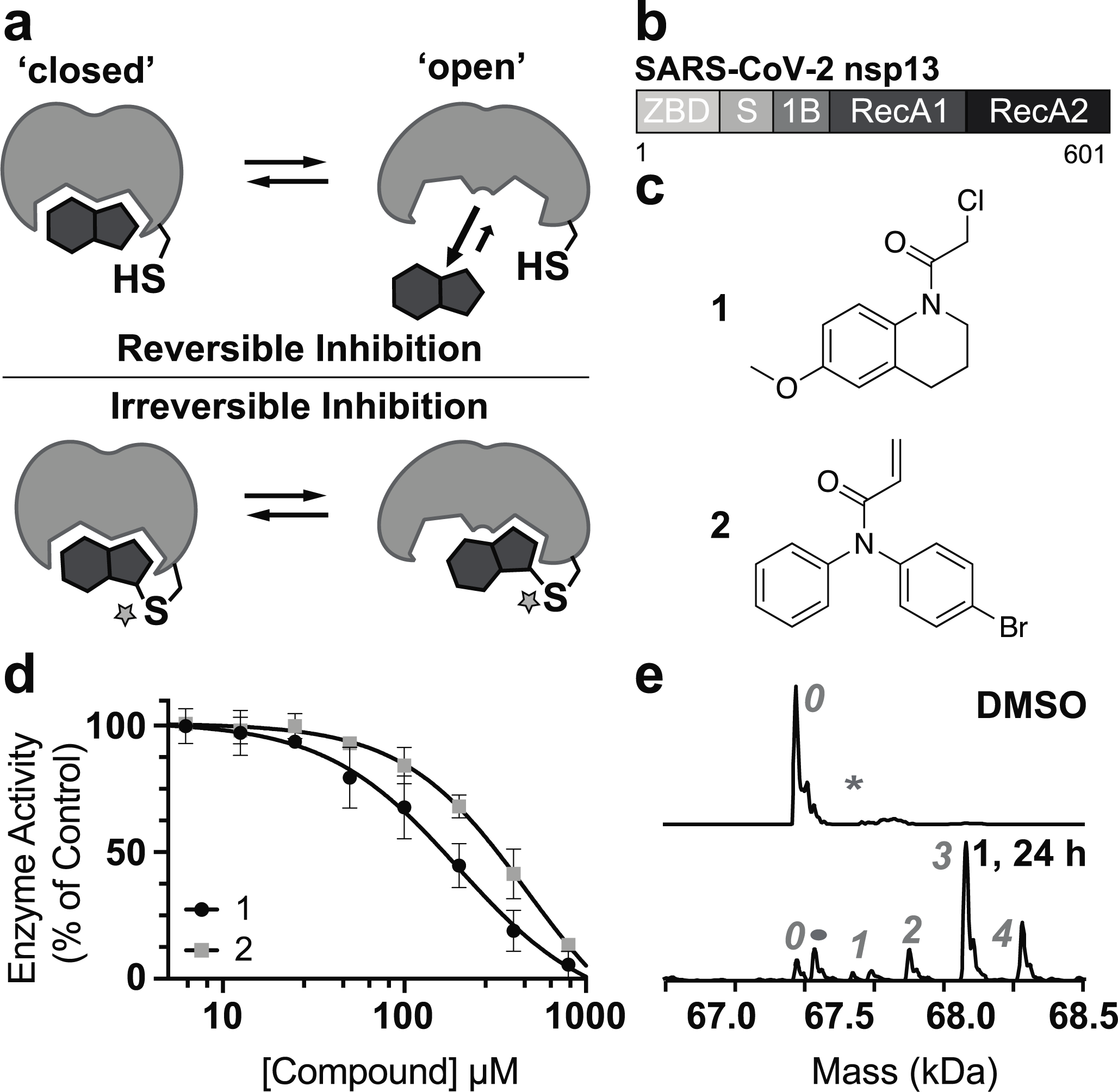
A ‘scout fragment’-based function-first approach to discover covalent inhibitors of helicases. (a) Schematic of irreversible verses reversible inhibition of a conformationally dynamic enzyme. (b) Domain organization of nsp13, a superfamily-1 helicase. (c) Chemical structure of ‘scout fragments’, compounds **1** (or KB02) and **2 (**or KB05). (d) Dose-dependent inhibition of nsp13 helicase activity by compounds **1** and **2** (IC_50_: compound **1** = 199 ± 107 μM, compound **2** = 475 ± 344 μM; 8h incubation; 4°C; n=2, mean ± SD). (e) Native MS analysis of nsp13 liganding by compound **1** (200 µM compound **1,** 24h incubation, 4°C; number of adducts: gray).

Our efforts were inspired by the use of electrophilic ‘scout fragments’ in large-scale chemical proteomic workflows profiling ligandability of thousands of nucleophilic residues (e.g. cysteine) across native proteomes^8–11^. While these studies enable proteome-wide identification of ligandable sites for covalent modification, methods to assess if ligand engagement modulates protein function are only recently emerging^12,13^. Chemical proteomics analyses with ‘scout fragments’ has identified ligandable sites in helicases^11^, however, it is unclear if any of these liganding events inhibit helicase activity and if these fragments can be progressed into site-specific inhibitors.

Here, we combine the use of electrophilic ‘scout fragments’ with biochemical assays, enantiomeric probe pairs, and mass spectrometry to identify site-specific helicase inhibitors. To develop our approach, we first focused on nsp13, the SARS-CoV-2 helicase, a member of the SF1 helicases that is required for SARS-CoV-2 replication^14,15^. Nsp13 has been proposed to be an important target of antiviral therapies due to its high degree of conservation across coronaviruses^16^. This enzyme consists of five distinct domains, including a zinc binding region (ZBD), a stalk (S) that connects to a 1B beta barrel domain, and two RecA domains, and can unwind DNA and RNA substrates (Figure 1B)^17–19^. Efforts to identify inhibitors of nsp13 have been reported^18,20–26^. However, it is unclear if hits identified from crystallographic fragment screens can inhibit helicase function^18^. In addition, target specificity has not been firmly established for inhibitors from high-throughput screening and drug repurposing efforts^20–22^. These compounds include suramin, an antiparasitic drug that has been shown to be a promiscuous binder of additional helicases^27^ and other SARS-CoV-2 proteins^28^. To our knowledge, direct-binding selective inhibitors of nsp13 have not been described.

## Results

To identify covalent inhibitors for nsp13 we selected two ‘scout fragments’ (compound **1**: KB02 and compound **2**: KB05, Figure 1C) that have been extensively characterized in chemical proteomic studies^8–11^. We generated recombinant nsp13 and adapted a fluorescence-based DNA unwinding assay for analysis of helicase activity^19^. This biochemical assay revealed dose-dependent inhibition of nsp13 by compounds **1** and **2** (Figure 1D). We selected the more potent compound **1** for subsequent studies.

Next, we used high-resolution native mass spectrometry (nMS) to measure the masses of intact protein and protein ligand complexes, establish ligand binding modes (covalent and/or noncovalent) and determine relevant stoichiometries. We observed predominantly three covalent adducts to nsp13 in the presence of compound **1** (200 μM, 24h incubation, Figure 1E). Nsp13 contains 26 cysteine residues^29^, and these data indicate that a subset of these are being liganded by compound **1**.

We next examined the liganding of nsp13 using site-mapping MS, a bottom-up proteomics method complementary to nMS. This approach indicated that compound **1** modifies C441 and C444, residues that are in a loop within nsp13’s ATP-binding pocket (Figure S1, Table S1). We next generated a construct with a double mutation (C441S C444S, hereafter nsp13^C441S C444S^), determined it to be enzymatically active (∼1.9-fold lower catalytic efficiency (k_cat_/K_m, dsDNA_) compared to nsp13^wt^; Figure S2, Table S2), and found that compound **1** still inhibited its helicase activity (Figure 2A).

**Figure 2.**
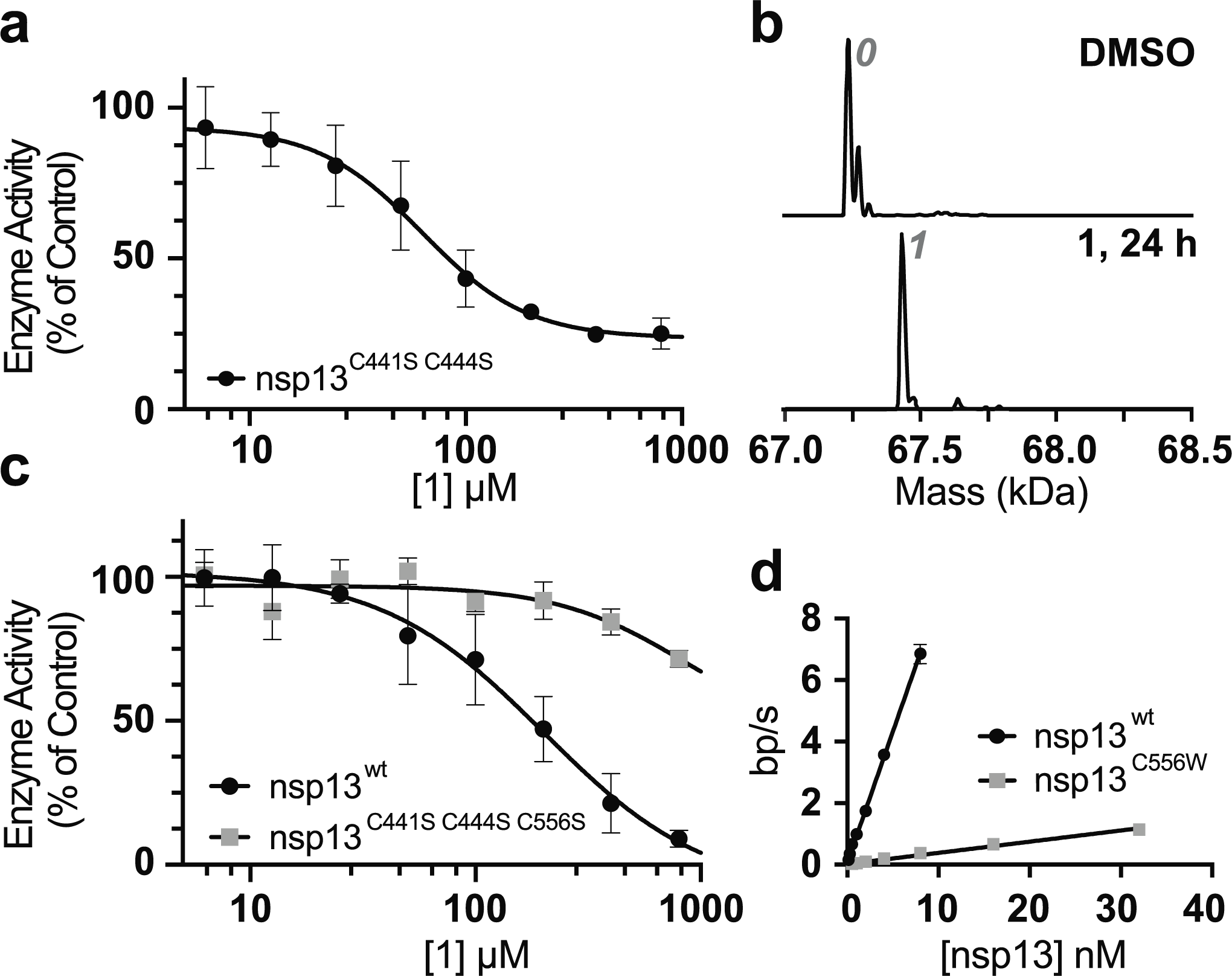
Identifying an allosteric site that can be liganded to inhibit nsp13 activity. (a) Dose-dependent inhibition of nsp13^C441S C444S^ helicase activity by compound **1** (IC_50_ = 62 ± 19 μM, 8h incubation, 4°C; n=2, mean ± SD). (b) Native MS analysis of nsp13^C441S C444S^ liganding by compound **1** (200 µM compound **1**, 24h incubation, 4°C.) (c) Dose-dependent inhibition of nsp13^wt^ and nsp13^C441S C444S C556S^ by compound **1** (8h incubation, 4°C). (d) Enzyme velocity (base pairs unwound per second; n=2, mean ± SD) versus concentration of nsp13^wt^ and nsp13^C556W^.

We performed nMS analysis on nsp13^C441S C444S^ and interestingly, only a single adduct was detected after incubation with compound **1** (200 μM, 24h, Figure 2B). This is consistent with the three covalent adducts observed when nsp13^wt^ was incubated with compound **1** (Figure 1E). Site-mapping MS experiments indicated that C556 in nsp13^C441S C444S^ is the predominant site of liganding by compound **1** (Figure S3, Table S3). We next generated a construct with a triple mutation (C441S C444S C556S, hereafter nsp13^C441S C444S C556S^), determined it to be enzymatically active (∼4 fold lower catalytic efficiency compared to nsp13^wt^; Figure S2, Table S2), and found that compound **1** does not substantially inhibit the helicase activity of nsp13^C441S C444S C556S^ (Figure 2C).

To test if liganding C556 would be sufficient to inhibit nsp13 helicase activity, we generated a nsp13 construct with a C556W (hereafter, nsp13^C556W^) point mutation, as it has been proposed that tryptophan mutations can mimic liganding by small molecules^30^. Enzymatic analysis suggests that the tryptophan mutant has a K_m, dsDNA_ ∼3 fold higher than nsp13^wt^ (K_m, dsDNA_: nsp13^wt^ = 573 ±1061 nM, nsp13^C556W^ = 1493 ± 642 nM; n=2 mean ± SD) (Figure S2, Table S2). However, the rate of unwinding is ∼20 fold less for nsp13^C556W^ compared to nsp13^wt^ (Figure 2D, Table S2). Taken together, these data suggest that covalent modification of C556, would suppress helicase activity.

Encouraged by these data we synthesized four analogs of compound **1** with simple modifications, such as methyl substitutions (Figure S4). Testing these compounds *in vitro* revealed that compound **3** was more potent than compound **1** (Figure 3A, 3B; Figure S4). We purified enantiomers of **3**, yielding compounds **3a** and **3b**, and found differences in the potency of nsp13 inhibition (Figure 3B). We next generated a computational covalent docking model focusing on the C556 residue (Figure S5). These analyses suggested that substitutions of compounds **3a/3b** could be designed to interact with a hydrophobic pocket proximal to C556 (Figure S5). We synthesized 7 additional analogs of compound 3 and found that aryl substituted compounds were substantially better inhibitors (Figure S6). Enantiomers of compound **4**, the most potent of these analogs, were purified to obtain compounds **4a** and **4b** (Figure 3C). Again, we observed differences in the potency of nsp13 inhibition based on the stereochemistry of the methyl group (Figure 3D). Interestingly, the stereochemistry of the more active analogs (**3b** and **4b**) is different. Nonetheless, the observed potency differences suggest that non-covalent contacts play a role in the ligand-enzyme interactions. Native MS analyses indicated that compound **4b**, the more potent enantiomer of compound **4**, ligands nsp13 predominantly at a single residue (Figure 3E).

**Figure 3.**
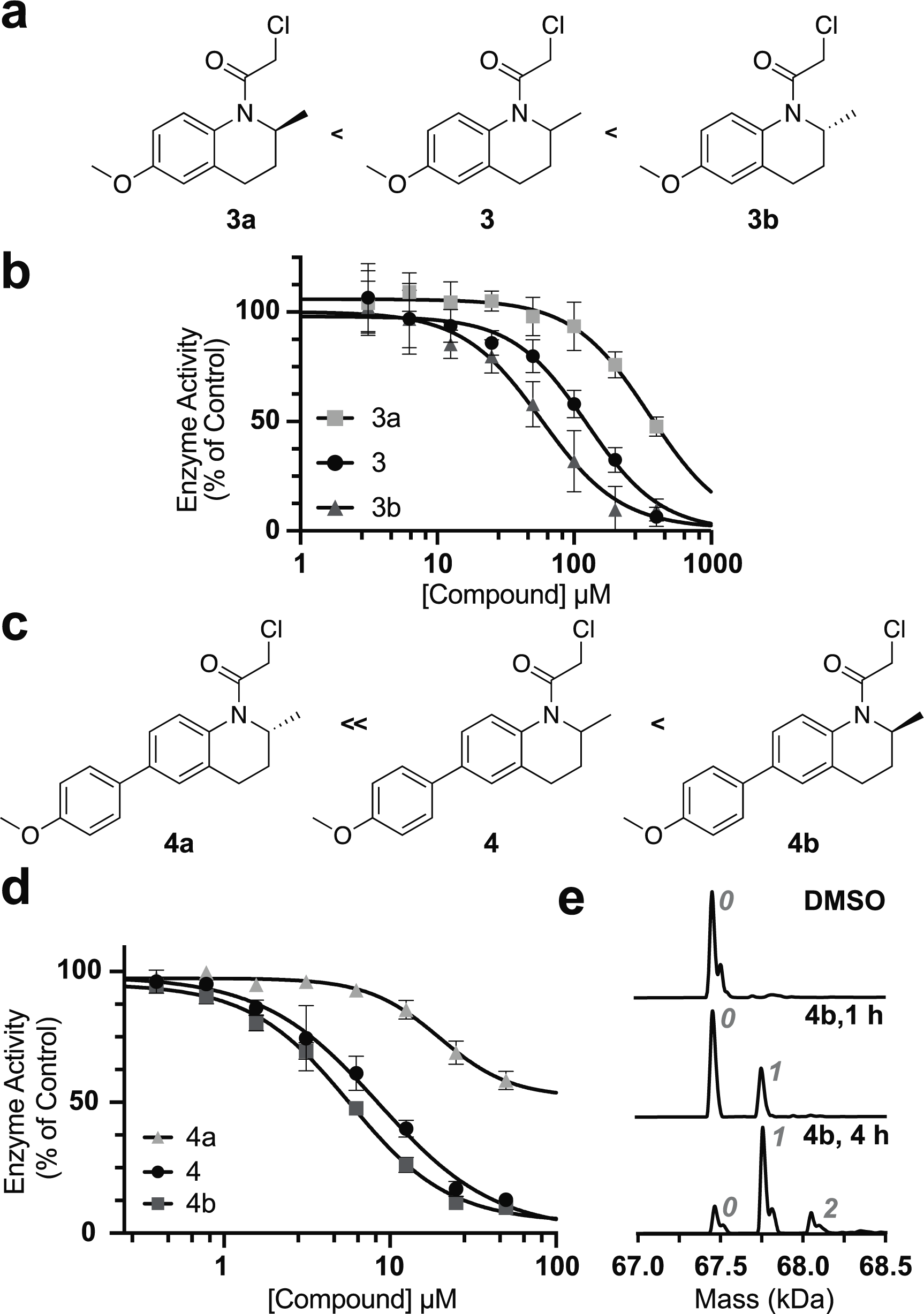
Characterizing analogs of compound **1**. (a) Chemical structures of compounds **3**, **3a** and **3b** and potency rank ordering (left to right, indicated by chevron). (b) Dose-dependent inhibition of nsp13 helicase activity by compounds **3**, **3a** and **3b** (IC_50_: compound **3a** ∼ 400 µM; compound **3** = 62 ± 29 µM; compound **3b** = 29 ± 9.4 µM; 8h incubation, 4°C; n=2, mean ± SD). (c) Chemical structures of compounds **4**, **4a** and **4b** and potency rank ordering (left to right, indicated by chevron). (d) Dose-dependent inhibition of nsp13 helicase activity by compounds **4**, **4a** and **4b** (IC_50_: compound **4a** ∼ 50 µM; compound **4** = 8.5 µM ± 2.4; compound **4b** = 5.7 ± 0.6 µM; 4-hour incubation, 4°C; n=2, mean ± SD). (e) Native MS analysis of nsp13 liganding by compound **4b** (20 µM compound **4b,** incubation times as indicated, 4°C)

Site-mapping MS analyses revealed C556 is the primary site of nsp13 liganding by compound **4b** (Figure S7, Table S4). We generated a construct with a point mutation at C556 (nsp13^C556S^), determined it to be enzymatically active (∼3 fold lower catalytic efficiency compared to nsp13^wt^; Figure S2, Table S2), and gratifyingly, we did not detect inhibition of this mutant construct by compound **4b** (Figure 4A). We also performed nMS experiments and did not detect modification of nsp13^C556S^ in the presence of compound **4b** (Figure 4B), indicating that this compound acts by selective covalent modification of the C556 residue in nsp13.

**Figure 4.**
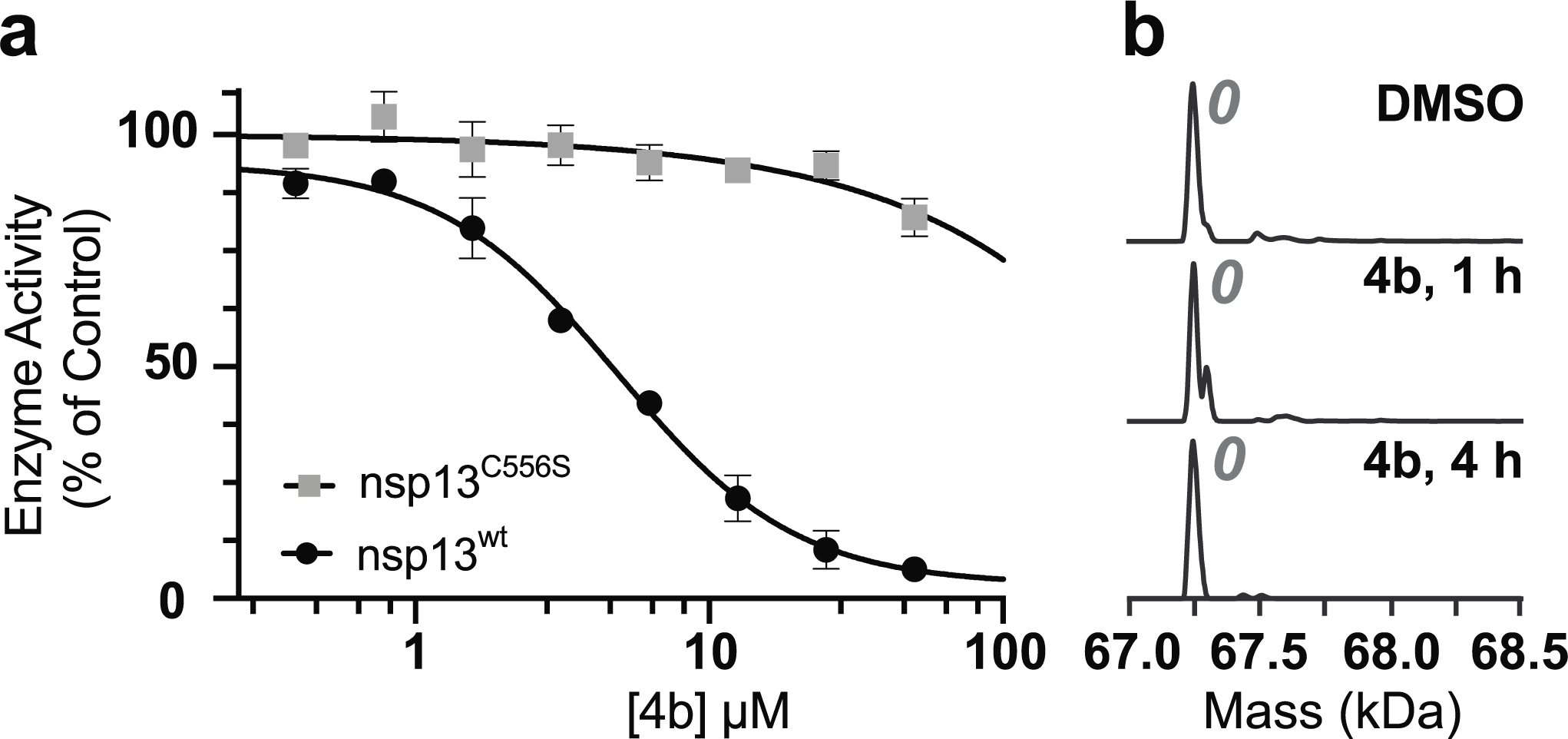
Site-selective liganding of nsp13 by compound **4b.** (a) Dose-dependent inhibition of nsp13^wt^ and nsp13^C556S^ by compound **4b** (IC_50_: nsp13^wt^ = 4.8 ± 0.6 µM, nsp13^C556S^ = N/A; 4h incubation; n=2, mean ± SD). (b) Native MS analysis of nsp13^C556S^ liganding by compound **4b** (20 µM compound **4b,** incubation times as indicated, 4°C).

To profile selectivity, we next tested inhibition of two mammalian helicases by compound **4b**. We selected the superfamily-2 RecQ helicases Bloom syndrome (BLM) and Werner syndrome (WRN), two enzymes involved in maintaining genome stability^31^. Importantly, WRN helicase has been identified as a unique vulnerability in certain cancer cell types^32–35^. Guided by literature precedent, constructs for BLM and WRN (BLM^636–1298^, WRN^515–1233^) were expressed in recombinant form and fluorescence-based activity assays were adapted (Figure S8)^36,37^. Each construct contains a helicase core, RecQ domain, and HRDC (helicase and RNAaseD c-terminal domain) domain (Figure 5A). We found that nsp13 is inhibited by compound **4b** more potently than WRN or BLM (Figure 5B). Together, these data indicate that compound **4b** is a selective, site-specific, allosteric inhibitor of nsp13.

**Figure 5.**
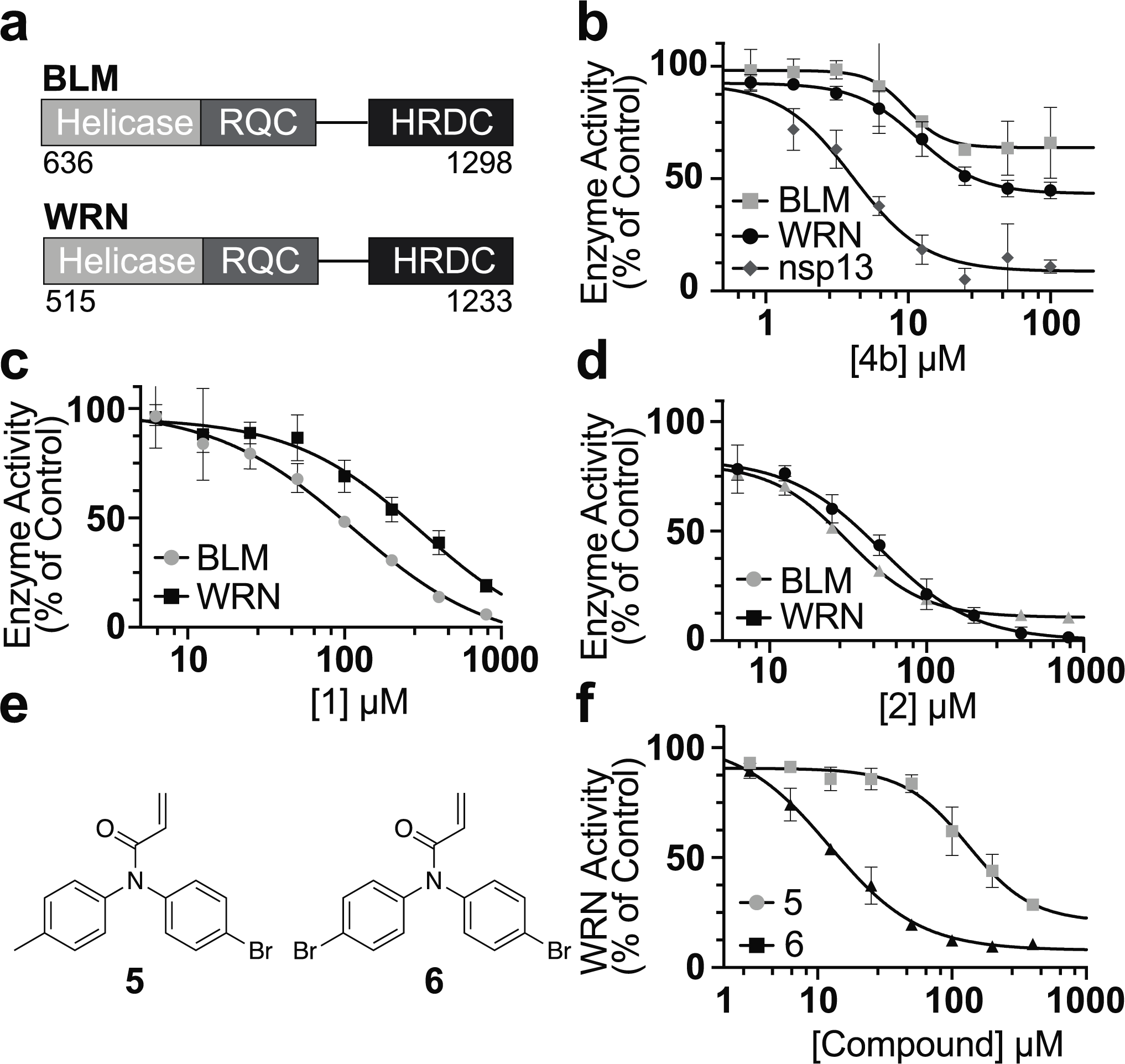
Characterizing inhibition of the helicases BLM and WRN by scout fragments and their analogs. (a) Domain organization of BLM and WRN constructs used. (b) Dose-dependent inhibition of nsp13, BLM and WRN helicase activity by compound **4b** (IC_50_: nsp13 = 4.1 ± 0.6 µM, BLM ∼ 50 µM, WRN ∼ 25 µM; 4-hour incubation, 4°C; n=2, mean ± SD). (c) Dose-dependent inhibition of BLM and WRN helicase activity by compound **1** (IC_50_: BLM = 308.0 ± 234.0 μM; WRN = 112.7 ± 21.65 μM; 8-hour incubation, 4°C; n=2, mean ± SD). (d) Dose-dependent inhibition of BLM and WRN helicase activity by compound **2** (IC_50_: BLM = 51.4 ± 4.9 μM; WRN= 31.3 ± 4.0 μM; 8h incubation, 4°C). (e) Chemical structures of compounds 5 and 6. (f) Dose-dependent inhibition of WRN by compounds **5** and **6** (IC_50_: compound **5** = 131.0 ± 28.8 µM, compound **6** = 12.2 ± 2.1 µM; 8-hour incubation, 4°C; n=2, mean ± SD)

We next examined if a function-first ‘scout-fragment’-based approach can be used to identify inhibitors of mammalian RecQ helicases. Chemical proteomic studies have suggested that both BLM and WRN can be partially liganded by compounds **1** and **2**^10,11^. We found that both BLM and WRN activity is weakly inhibited by compound **1**, but more strongly inhibited by compound **2** (Figure 5C, 5D). Time-dependent inhibition suggests a covalent mode of action (Figure S8). We synthesized and tested two analogs of compound **2**, yielding compounds **5** and **6** (Figure 5E), and found that compound **6** is a more potent inhibitor of WRN than compound **2** (Figure 5F). These data suggest that compounds **2** and **6** could be useful starting points to develop covalent inhibitors for RecQ helicases.

## Discussion

Together, our findings suggest a function-first approach, based on biochemical testing of electrophilic scout fragments combined with the use of enantiomeric probe pairs and mass spectrometry, to identify selective covalent inhibitors for helicase mechanoenzymes. In the case of nsp13, of the 26 cysteines, we identified a single residue in an allosteric site that is liganded by compound **4b** to inhibit helicase activity. Structural models reveal that C556 is >10 Å from nsp13’s RNA-binding channel and is not likely to be directly involved in ATP- or RNA-binding^38^. Further studies will be required to understand how liganding C556 in nsp13 allosterically inhibits helicase activity. Our finding that nsp13 activity is reduced by mutating this cysteine to serine (Figure S2, Table S2), a conservative change in this amino acid, suggests that potential inhibitor resistance-conferring mutations would have an associated fitness cost for the virus. Consistent with this hypothesis, sequences of SARS-CoV-2 submitted to the GISAID database indicate a low frequency of C556 mutations (132 of >15 million sequences available)^39^.

For the few helicases for which selective chemical inhibitors have been reported^40–44^, targeting allosteric sites has been an effective strategy. In the case of the hepatitis C NS3 helicase integrative approaches, which included high-throughput screens, fragment screens and structural analysis, were employed to identify druggable allosteric sites^40,41^. Both allosteric and orthosteric inhibitors of the Brr2 helicase, an enzyme essential for spliceosome function, were identified and the allosteric binders exhibited better specificity^42^. Characterizing the mechanisms of action of natural products (e.g. rocaglamids) or screening hits has also serendipitously identified druggable allosteric sites in EIF4A^43^ and BLM helicases^44^, respectively. Our data suggest that characterizing scout fragment-based activity inhibition may present a robust approach to identify chemical inhibitors that selectively target allosteric sites in helicases and other conformationally dynamic mechanoenzymes.

## Associated Content

The Supporting Information is available as an associated PDF

## ACKNOWLEDGEMNTS

T.M.K. is grateful to the NIH/NIGMS (GM98579 and R35GM130234-01) and the Stavros Niarchos Foundation for funding. E.V.V is grateful to The Rockefeller University start-up funds, The Robertson Foundation, and the Achelis and Bodman Foundation for funding. E.V.V. is also grateful to the Searle Scholars Program. J.R.R., L.E.V and H.S were supported by the Tri-Institutional Ph.D. Program in Chemical Biology and The Rockefeller University Graduate program. L.E.V and H.S were also supported by the NIH T32 (GM115327 and GM136640) Chemistry-Biology Interface Training Grant to the Tri-Institutional Ph.D Program in Chemical Biology. B.T.C is grateful to the NIH for funding (P41 GM109824 and P41 GM103314). Support for this project was also provided by the Sanders Tri-Institutional Therapeutics Discovery Institute (TDI), a 501(c) (3) organization. TDI receives financial support from Takeda Pharmaceutical Company, TDI’s parent institutes (Memorial Sloan Kettering Cancer Center, The Rockefeller University and Weill Cornell Medicine) and from a generous contribution from Lewis Sanders and other philanthropic sources. We thank Caroline Webster for assistance with site-mapping MS experiments.

## General Methods

### Vectors for Recombinant Protein Expression

Vector for expression of SARS-CoV-2 wild-type nsp13 was obtained as previously described^1^. Briefly, the sequence for full length nsp13 was cloned into a pet28(a)+ vector by Genescript, after which an HRV-3C (PreScission) protease cleavage site was introduced using a Quikchange mutagenesis (Agilent). This construct was used for all experiments with nsp13^wt^ (enzymatic assays, nMS, and site-mapping MS).

Vectors for the expression of mutant SARS-CoV-2 nsp13 variants C441S C444S, C441S C444S C556S, C556W, and C556S were generated by performing Quikchange mutagenesis (Agilent) on a previously described plasmid purchased from Addgene (#159614)^2^. Briefly, the whole vector was PCR amplified using overlapping primers that contained the mutation of interest. Vector amplification was verified by 1% agarose gel, and successful PCR reactions were transformed directly into DH5*α E. Coli.* cells.

Vector for the expression of human wild-type Bloom’s syndrome helicase (Hs-BLM-636-1298) was based on previously described constructs^3,4^. The vector was generated by cloning cDNA (Dharmacon) into a bacterial expression vector with an N-terminal His6 tag and HRV 3C (PreScission) protease cleavage site.

Vector for the expression of human wild-type Werner’s syndrome helicase (Hs-WRN-515-1233) was generated by cloning codon optimized cDNA (Genscript) into a pFastBac backbone for insect cell expression with an N-terminal His6 tag and TEV cleavage site.

Sequence verification for all vectors was carried out by nanopore sequencing (Plasmidsaurus). Sequence alignments were done using SnapGene pro.

### Protein Expression and Purification

Protocol for the expression and purification of wild type SARS-CoV-2 nsp13 was performed as previously described^1^.

Protocol for expression and purification of mutant SARS-CoV-2 nsp13 variants was adapted from Newman et. al^2^. Briefly, the plasmid was transformed into BL21 Rosetta cells and cultures were grown in Miller’s LB medium (LMM, Formedium) at 37°C, until OD_600_ reached 0.7. The cultures were chilled to 18°C, expression was induced by the addition of 200 μM IPTG (GoldBio), and the cultures allowed to shake until the next morning (∼18-20 hours). The next day, bacteria were harvested by spinning down at 5000 g for 10 minutes. Cell pellets were resuspended in lysis buffer (50 mM HEPES pH 7.5, 500 mM NaCl, 5% Glycerol, 0.1 mM PMSF, 0.5 mM TCEP, 1 c0mplete protease tab (50 mL buffer + tablet; Roche)). Lysis was carried out either by probe sonication (7 minutes, 10 seconds on, 10 seconds off, 70% amplitude) or Emulsiflex-C5 homogenizer (Avestin, 5-7 cycles at 10-12 kPsi). Lysate was then cleared using a Beckman Coulter ultracentrifuge and a Type 45 Ti rotor at 30,000 rpm for 30 minutes at 4 °C. The supernatant was then mixed with Ni-NTA resin (Qiagen) and allowed to incubate at 4 °C with rocking for 1 hr. The supernatant and resin mixture was then passed through a gravity column, and the resin was subsequently washed with ∼40 mL of lysis buffer. The resin was then washed with 20 mL lysis buffer + 45 mM imidazole, 10 mL Hi-salt buffer (lysis buffer + 1M NaCl), and 10 mL lysis buffer + 45 mM imidazole. Once complete, the protein was eluted in batch with 20 mL lysis buffer + 300 mM imidazole (Ni-NTA elution buffer). The elution was loaded onto a High Trap SP HP column (Cytiva) at a flow rate of 1 mL/min, followed by washing the column with 10 mL Ni-NTA elution buffer. The protein was eluted from the column over 10 mL Hi-salt buffer. TEV protease was added to the combined elution fractions and the protein was left to dialyze overnight in dialysis buffer (50 mM HEPES, 1000 mM NaCl, 5% Glycerol, 1 mM TCEP). The next day, 1 mL of Ni-NTA beads were equilibrated with reverse nickel buffer (50 mM HEPES, 500 mM NaCl, 30 mM Imidazole, 5% Glycerol, 1 mM TCEP). Cleaved nsp13 was supplemented with 45 mM imidazole, sample was passed over Ni-NTA beads and flowthrough was collected. Protein was then injected onto a Superdex 200 increase gel filtration column which was equilibrated with gel filtration buffer (50 mM HEPES pH 7.5, 500 mM NaCl, 0.5 mM TCEP). Fractions containing protein were checked by SDS-PAGE, then peak fractions were pooled, aliquoted and flash frozen in gel filtration buffer.

Protocol for the expression and purification of BLM^636–1298^ was adapted from Nguyen et. al^3^. and Newman et. al^4^. Briefly, the plasmid was transformed into BL21 Rosetta cells and cultures were grown in Miller’s LB medium (LMM, Formedium) at 37°C, until OD_600_ reached 1.8. The cultures were chilled to 18°C, expression was induced by the addition of 200 μM IPTG (GoldBio), and the cultures allowed to shake until the next morning (∼18-20 hours). The next day, bacteria were harvested by spinning down at 5000 g for 15 minutes. Cell pellets were resuspended in lysis buffer (50 mM HEPES·Na pH 7.6, 500 mM NaCl, 5% glycerol, 1 mM TCEP) and supplemented with 1 mM PMSF, 1 cOmplete protease inhibitor tablet (50 mL buffer + tablet; Roche), and 5 uL benzonase. Lysis was carried out either using an Emulsiflex-C5 homogenizer (Avestin, 5-7 cycles at 10-12 kPsi). Lysate was then cleared using a Beckman Coulter ultracentrifuge and a Type 45 Ti rotor at 30,000 rpm for 30 minutes at 4 °C. The supernatant was then mixed with Ni-NTA resin (Qiagen) and allowed to incubate at 4 °C with rocking for 1 hr. The supernatant and resin mixture was then passed through a gravity column, and the resin was subsequently washed with 250 mL (50 mM HEPES·Na pH 7.6, 500 mM NaCl, 5% glycerol, 1 mM TCEP, 20 mM imidazole). Elution buffer (50 mM HEPES·Na pH 7.6, 500 mM NaCl, 5% glycerol, 1 mM TCEP, 300 mM imidazole) was then added to the bead slurry and eluted until no protein was detectable by Bradford reagent. The elution was loaded onto a heparin column (Cytiva) at 1 mL/min, followed by washing the column with 30 mL heparin A buffer (50 mM HEPES·Na pH 7.6, 150 mM NaCl, 5% glycerol). The protein was eluted over a linear gradient of 0-100% heparin A/heparin B buffer (50 mM HEPES·Na pH 7.6, 1 M NaCl, 5% glycerol). Pooled fractions were incubated overnight at 4°C with PreScission protease. The next day, 1 mL of Ni-NTA beads were equilibrated with reverse nickel buffer (50 mM HEPES·Na pH 7.6, 500 mM NaCl, 5% glycerol, 30 mM imidazole). Cleaved BLM^636–1298^ was supplemented with 30 mM imidazole, sample was passed over Ni-NTA beads and flowthrough was collected. Protein was then injected onto a Superdex 200 increase gel filtration column which was equilibrated with gel filtration buffer (50 mM HEPES·Na pH 7.5, 500 mM NaCl, 5% glycerol). Fractions containing protein were checked by SDS-PAGE, then peak fractions were pooled, aliquoted and flash frozen in gel filtration buffer.

WRN^515–1233^ was expressed and purified from insect cells. Briefly, recombinant baculovirus was generated using the Bac-to-Bac system (ThermoFisher). Hi5 cells (3 x 10^6^ cells/mL) were infected with P3 virus (50/1 v/v) and allowed to incubate at 27°C for 72 hours shaking at 115 rpm. Cells were harvested by centrifugation at 3000 g for 10 minutes at 4°C, after this, the cell pellets were resuspended in lysis buffer (50 mM Tris-HCl pH 8.0, 500 mM NaCl, 20 mM imidazole, 2 mM MgCl_2_, 5% glycerol, 2.5 mM bME) supplemented with 1 mM PMSF and cOmplete protease inhibitor cocktail (50 mL buffer + tablet; Roche). Lysis was performed with a Dounce homogenizer (Thomas Scientific), 2 x 20 strokes for each volume of cells. Lysate was clarified by centrifugation at 45,000 rpm for 45 min at 4°C (Type 70 Ti rotor) followed by filtration with a 0.22 µm filter. Clarified lysate was then incubated with equilibrated Ni-NTA beads (Qiagen) for 1 hour at 4°C with nutation. Bead slurry was applied to a gravity column, washed with 200 mL lysis buffer, and eluted with the minimal amount of elution buffer (50 mM Tris-HCl pH 8.0, 500 mM NaCl, 300 mM imidazole, 2 mM MgCl_2_, 5% glycerol, 2.5 mM bME), fractions were checked for protein content using Bradford reagent. Those with the highest protein concentration were pooled and TEV protease was added, the mixture was then dialyzed for 3 hours at 4°C in dialysis buffer (50 mM Tris-HCl pH 8.0, 150 mM NaCl, 20 mM imidazole, 2 mM MgCl_2_, 5% glycerol, 2.5 mM bME). Dialyzed sample was loaded onto equilibrated Ni-NTA resin, flow through was collected and passed over the bead slurry twice more. A small volume reverse Ni-NTA buffer (50 mM Tris-HCl pH 8.0, 150 mM NaCl, 20 mM imidazole, 2 mM MgCl_2_, 2.5 mM bME) was added to elute any remaining protein, content was checked by Bradford reagent. Protein was then loaded onto a heparin column (Cytiva) equilibrated in heparin A buffer (50 mM Tris-HCl pH 8.0, 150 mM NaCl, 2 mM MgCl_2_, 5 mM bME). Elution was performed using a linear gradient of 0-100% heparin B buffer (50 mM Tris-HCl pH 8.0, 1 M NaCl, 2 mM MgCl_2_, 5 mM bME) over 25 minutes. Fractions containing protein were checked using SDS-PAGE, and the peak fractions were pooled. Heparin eluate was then loaded onto a Superdex200 Increase 10/300 column equilibrated in gel filtration buffer (25 mM HEPES-Na pH 7.5, 300 mM NaCl, 2 mM MgCl_2_, 1 mM TCEP). Fractions containing protein were checked by SDS-PAGE, then peak fractions were pooled, aliquoted and flash frozen in gel filtration buffer.

### DNA Substrate Preparation

Preparation of DNA duplex substrate for nsp13 was adapted from Mickolajcyzk et. al^5^. Briefly, oligonucleotides were purchased from Integrated DNA technologies with fluorescent and quenching labels. Oligos were mixed in a 1.2:1 ratio of quencher (5’ - TTTTTTTTTTCTGATGTTAGCAGCTTCGT-TAMRA-3’) to fluorescent strand (5’ -IAbRQ-ACGAAGCTGCTAACATCAG -3’), supplemented with 20 mM Tris pH 7.6 and 50 mM KCl, and annealed by heating to 95°C, followed by cooling at a rate of 1°C/minute down to 25°C.

For use in nsp13 helicase assays, an un-labelled “capture” oligo with the sequence 5’-CTGATGTTAGCAGCTTCGT-3’ was ordered from Integrated DNA Technologies and used.

Short-forked DNA (sfDNA) substrate for BLM and WRN was adapted from Newman et. al^4^. Briefly, oligonucleotides were purchased from Integrated DNA technologies with fluorescent and quenching labels. Oligos were mixed in a 1.2:1 ratio of quencher (5’ - TTTTTTAGCGTCGAGATC-BHQ2-3’) to fluorescent strand (5’ -TAMRA-GATCTCGACGCTCTCCCCTCCC -3’), supplemented with 20 mM Tris pH 7.6 and 50 mM KCl, and annealed by heating to 95°C, followed by cooling at a rate of 1°C/minute down to 25°C.

For use in BLM and WRN helicase assays, an un-labelled “capture” oligo with the sequence 5’-GATCTCGACGCT-3’ was ordered from Integrated DNA Technologies and used.

### Analysis of Helicase Activity

All assays for analysis of nsp13, BLM and WRN helicase activity were performed in black 384 well plates (Corning) with a final assay volume of 24 μL. Assays were initiated by the addition of ATP and fluorescence was measured every 15 seconds for 30 minutes using a Synergy NEO 2 Microplate reader (λ excitation = 544 nm, λ emission = 590 nm). Fluorescence values were plotted against time and initial velocity was calculated by determining the slope over the first ∼300 seconds. To correct for background signal, unwinding calculated from the no ATP control was subtracted from all experimental wells. Using a fluorescent oligo calibration curve, RFU values were converted into nM substrate unwound. For inhibition assays, percentage was calculated by normalizing values to the DMSO control.

All assays with nsp13 were performed in nsp13 assay buffer (20 mM HEPES (pH 7.5), 40 mM KCl, 5 mM MgCl2, 0.5 mM EDTA, 2.5 mM glutathione, 0.01% Triton X-100, and 0.1 mg/mL bovine serum albumin). Helicase assays to determine nsp13 enzymatic parameters K_M_ and k_cat_ were performed by mixing nsp13 (0.5-2 nM (16 nM for nsp13^C556W^; in assay buffer) in a 2x solution with labelled duplex DNA and capture DNA (0-2000 nM labelled duplex DNA, 2000 nM capture DNA) in a 4x solution. Assays were initiated by the addition of ATP as a 4x stock (150 mM final).

Helicase inhibition assays were performed by mixing a 4x nsp13 solution (0.5-2 nM; in assay buffer) with 2x solutions of inhibitor (in assay buffer, 1-4% DMSO final). Assay plate was then allowed to incubate at 4°C for the designated amount of time. At the given time point, a 4x solution of duplex DNA and capture DNA (100 nM duplex DNA, 400 nM capture DNA) was added. Assays were then initiated by addition of a 4x ATP (150 mM) solution.

BLM and WRN helicase inhibition assays were performed by mixing a 4x BLM/WRN solution (10 nM BLM, 15 nM WRN; in assay buffer) with 2x solutions of inhibitor (in assay buffer, 1-4% DMSO final). Assay plate was then allowed to incubate at 4°C for the designated amount of time. At the designated time point, a 4x solution of duplex DNA and capture DNA (250 or 400 nM short-forked DNA, 1000 or 1600 nM capture DNA respectively) was added. Assays were then initiated by addition of a 4x ATP (1 mM) solution.

### Equations used for data fitting

The Michaelis-Menten equation was used to fit initial velocity values and calculate the parameters K_m_ and k_cat_ as a function of substrate concentration (Prism v.10.0 GraphPad Software).

IC_50_ values were calculated by plotting helicase activity relative to the positive control versus inhibitor concentration, and then fitting to sigmoidal equation (1). Values for at least two independent replicates were averaged and standard deviations were calculated (Prism v.10.0 GraphPad Software).

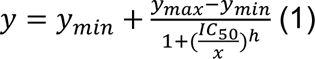

### Native Mass Spectrometry Analysis

#### Sample Preparation

2 µM of the wild-type or mutant nsp13 in assay buffer (20 mM HEPES pH 7.5, 40 mM KCl, 1 mM MgCl_2_ 2.5 mM GSH, 0.5 mM EDTA, 0.01% Triton X100, 2 % DMSO final) was incubated with 100-fold molar excess of compound **1** or 10-fold molar excess of compound **4b** at 4 °C. A 13-µL aliquot of each reaction mixture was taken at specific time points (t = 0, 1, 4, 24 h) and subsequently buffer-exchanged into nMS solution (150 mM ammonium acetate, 0.01% Tween-20) using Zeba desalting microspin columns with a 40-kDa molecular weight cut-off (Thermo Scientific). The buffer-exchange step also removed the excess unreacted compound. An aliquot (2 – 3 µL) of the buffer-exchanged sample was then loaded into a gold-coated quartz capillary tip that was prepared in-house and was electrosprayed into an Exactive Plus EMR instrument (Thermo Fisher Scientific) using a modified static nanospray source^6^.

#### nMS Analysis

The nMS parameters used included: spray voltage, 1.21 kV; capillary temperature, 150 or 200 °C; S-lens RF level, 200; resolving power, 8,750 or 17,500 at *m/z* of 200; AGC target, 1 x 10^6^; number of microscans, 5; maximum injection time, 200 ms; in-source dissociation (ISD), 10 V; injection flatapole, 8 V; interflatapole, 7 V; bent flatapole, 6 V; high energy collision dissociation (HCD), 125 – 150 V; ultrahigh vacuum pressure, 5.3 × 10^−10^ mbar; total number of scans, 100. Mass calibration in positive EMR mode was performed using cesium iodide. Raw nMS spectra were visualized using Thermo Xcalibur Qual Browser (version 4.2.47). Data processing and spectra deconvolution were performed using UniDec version 4.2.0 (Reid et. al., 2019; Marty et. al., 2015). The expected mass for the wild-type nsp13 is 67,464 Da (post-protease cleavage with three coordinated Zn^2+^ ions and nine deprotonated cysteine residues). The expected masses for the single and double Cys-to-Ser mutants are 67,244 and 67,228 Da, respectively. The predicted average masses for compounds **1** and **4b** were 239.7 Da and 329.8 Da, respectively. Reaction of each compound with a cysteine in nsp13 leads to the release of HCl (mass of +203 Da for compound **1** and +293 Da for compound **4b** for each covalently attached compound). The measured masses for the unreacted and modified proteins were within ± 2 Da from their respective expected masses.

### Site-Mapping Mass Spectrometry Analysis

#### Sample Preparation

Liganding reactions were assembled using 2 µM nsp13 and 200 µM compound **1** or 20 µM compound **4b** in assay buffer (20 mM HEPES (pH 7.5), 40 mM KCl, 1 mM MgCl_2_, 2.5 mM GSH, 2% DMSO final). Reactions were allowed to proceed for 24 hours for compound **1**, and 4 hours for compound **4b**. At the designated time point, reactions were quenched reaction with 200 µL ice cold acetone and chilled at −20°C for at least 30 minutes. Protein was pelleted by spinning at 16,000 rpm at 4°C, for 10 minutes. Supernatant was aspirated and sample was allowed to air dry. Pellet was resuspended in 50 µL resuspension buffer (9 M Urea, 50 mM TEAB, 10 mM DTT, made fresh) and sonicated for ∼5 minutes. Samples were then heated at 65°C for 15 minutes. 2.5 uL of 400 mM IA (20 mM final, made fresh) was added and samples were allowed to incubate for 30 minutes at 37 °C with shaking. Sample was diluted down to 1.8 M urea with 50 mM TEAB, then either 1 µg trypsin (for compound **1** preparations; 0.25 µg/uL, 4 uL; Promega) or 1 µg trypsin/LysC (for compound **4a/4b** preparations; 0.25 µg/uL, 4 uL total; Promega) was added and samples were allowed to incubate for 4 hours at 37 °C. For compound **1** preparations, this incubation proceeded overnight. For compound **4a/4b** preparations, sample was then diluted down to 0.9 M urea using 50 mM TEAB, and 1 µg GluC protease was added, samples were allowed to incubated overnight at 37 °C. The next morning, samples were acidified with formic acid and desalted with C-18 sep-pak columns using the following protocol: wet resin with 3 x 1 mL acetonitrile (ACN), equilibrate resin with 3 x 1 mL buffer A (95% H_2_O, 5% MeCN, 0.1% formic acid), slowly load sample onto column, collect flowthrough and pass over resin a second time, was with 3 x 1 mL buffer A, elute with 0.3 mL buffer B (80% MeCN, 20% H_2_O, 0.1% formic acid), collect and elute a second time. Elutions were then dried under reduced pressure and saved until MS analysis.

#### LC-MS/MS Analysis

Samples were analyzed by liquid chromatography tandem mass-spectrometry (LC-MS/MS) using an Orbitrap Eclipse mass spectrometer (Thermo Scientific) coupled to an UltiMate 3000 Series Rapid Separation LC system and autosampler (Thermo Scientific Dionex). The residue was re-dissolved in buffer containing 95% H_2_O, 5% CH_3_CN, and 0.1% FA (10 µL) and analyzed by LC-MS/MS. The peptides were loaded onto a capillary column (75 µm inner diameter fused silica, packed with C18 (PepMap^TM^, 100 Å, 2.0 µm, 25 cm) and separated at a flow rate of 0.25 µL/min using the following LC-MS gradient: 5% buffer B in buffer A from 0-10 min, 5-20% buffer B from 10-120 min, 20-45% buffer B from 120-140 min, 45-95% buffer B from 140-145 min, 95% buffer B from 145-147 min, 5% buffer B from 147-149 min, 95% buffer B from 149-151 min and 5% buffer B from 151-160 min (buffer A: 100% H_2_O, 0.1% FA; buffer B: 100% CH_3_CN, 0.1% FA). For peptide selection, fragmentation, and analysis, the scan sequence began with an MS1 master scan (Orbitrap analysis, resolution 240,000, 400−1600 m/z, RF lens 40%, charge state 2-7) with dynamic exclusion enabled (repeat count 1, duration 60 s). MS2 analysis consisted of quadrupole isolation (isolation window 0.7) of precursor ion followed by collision-induced dissociation (CID) in the ion trap with rapid scan rate (fixed collision energy 30% with 10 ms CID activation time, Activation Q = 0.25, Standard AGC, maximum injection time 100 ms). The MS2 files were extracted from the raw files using RAW Converter (version 1.1.0.22; available at http://fields.scripps.edu/rawconv/), uploaded to Integrated Proteomics Pipeline (IP2), and searched using the ProLuCID algorithm (publicly available at http://fields.scripps.edu/downloads.php) using a FASTA database containing SARS-CoV-2 P0DTD1 sequence, corresponding C441S mutant or C441S and C444S double mutant sequences from UniProt. Cysteine residues were searched with a static modification for carboxyamidomethylation (+57.02146 Da) and a differential modification containing peptide adducts with the compounds (+146.0732 Da for compound **1**, +236.1201 Da for Compound **4**). Peptides were required to be at least 5 amino acids long, and to have at least one tryptic or GluC terminus. ProLuCID data was filtered through DTASelect (version 2.0) to achieve a peptide false-positive rate below 1%.

### Covalent Docking Protocol

All modeling works were performed using the Schrödinger Suite (Release 2021-1, Maestro, Schrödinger, LLC, New York, NY). The Protein Preparation Wizard was used for the preparation of the SARS-Cov-2 helicase structure in complex with a fragment Z1429867185 (PDB 5RLE, chain B)^2^ with default settings. Ligand and water molecules were removed. The covalent docking workflow CovDock was applied for covalent docking using the Pose Prediction mode^7^. C556 was selected as the reactive residue. Compounds **3a** and **3b** were prepared using LigPrep and underwent “nucleophilic substitution” to yield the final poses. Then, the Sitemap tool^8^ was used to evaluate and characterize the binding site of a 6Å region around the ligand.

### Chemical Synthesis Information

Proton NMR spectra were recorded on Bruker Avance Neo spectrometer (600 MHz) with a 5 mm TXI probe or Bruker Avance DPX400 spectrometer operating at 400.13 MHz carrying a BBFO probe. Chemical shifts are reported as δ units of parts per million (ppm) relative to tetramethylsilane (δ 0.0). Multiplicities are reported: s (singlet), d (doublet), t (triplet), q (quartet), dd (doublet of doublets), m (multiplet) or br. (broad signal). Coupling constants are reported as a J value in Hertz (Hz). Solvents and reagents were purchased from Sigma-Aldrich. Commercially available reagents were used without further purification. Solvent evaporation was performed under reduced pressure using a rotary evaporator (Büchi). Reaction progress was monitored by thin layer chromatography or LC-MS (Waters Acquity H-Class UPLC/MS with QDa mass spectrometer using water + 0.1% formic acid (solvent A) and acetonitrile + 0.1% formic acid (solvent B). All flash chromatography was performed using standard silica gel unless specified otherwise.

## Synthetic Procedures

### General procedure A for hydrogenation

Unsaturated starting material (1.0 eq) was dissolved in toluene and Pd/C was added under an N_2_ atmosphere. The suspension was degassed, purged with H_2_ 3 times, and then stirred under H_2_ (50 psi). Reaction progress was monitored by TLC and LC/MS. Upon completion, the reaction was filtered and concentrated under reduced pressure. Crude products were used without further purification.

### General procedure B for chloroacetamide preparation

Amine starting material amine (1.0 eq) was dissolved in dichloromethane (DCM) or tetrahydrofuran (THF), and pyridine (3.0 eq) or triethylamine (TEA; 2.0 eq) was added. The mixture was stirred, cooled to 0 °C in an ice bath, and chloroacetyl chloride (1.0-3.0 eq) was added dropwise. Reactions were allowed to come up to room temperature and progress was monitored by TLC and LC-MS. Then the reactions were diluted with EtOAc (3 mL) and washed with sat. NaHCO_3_ (1 x 3 mL). The organic layers were combined, dried with MgSO_4_, filtered, and concentrated *in vacuo*. The crude residue was purified by flash chromatography or prep HPLC (Column: Waters Xbridge BEH C18 OBD Prep, specifications: 150×40mm×10um).

### General procedure C for acrylamide preparation

Commercially available amine starting material amine (1.0 eq) and dichloromethane (DCM), and pyridine (3.0 eq). The mixture was stirred, cooled in an ice bath, and acryloyl chloride (3.0 eq) was added dropwise. Reaction progress was monitored by TLC and LC-MS. Then the reactions were diluted with CH_2_Cl_2_ (3 mL) and washed with sat. NaHCO_3_ (1 x 3 mL). The organic layers were combined, dried with MgSO_4_, filtered, and concentrated *in vacuo*. The crude residue was purified by flash chromatography to afford pure product.

Compounds **1** and **2** were prepared according to literature precedence (Backus et. al.).

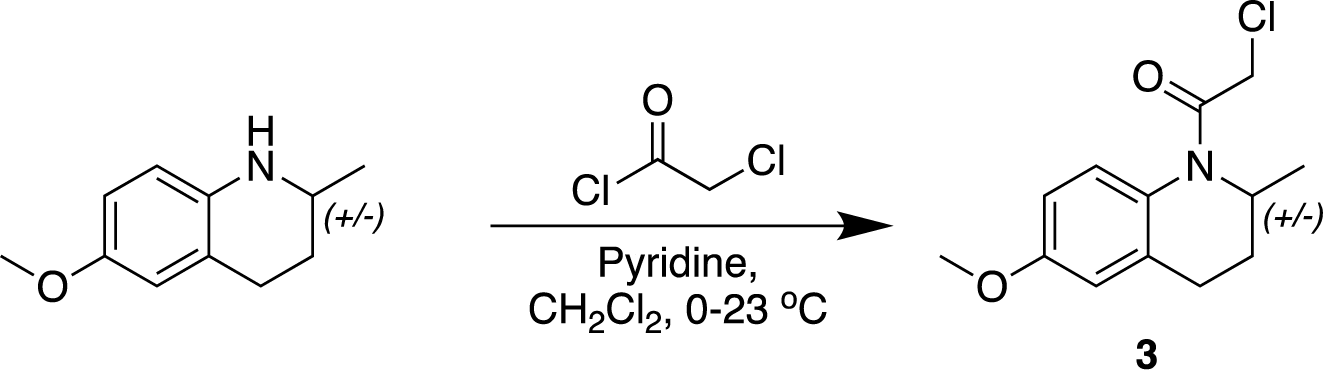

### 2-Chloro-1-(6-methoxy-2-methyl-3,4-dihydroquinolin-1(2H)-yl)ethan-1-one (3)

General procedure B was used with the following reagents: 6-methoxy-2-methyl-1,2,3,4-tetrahydroquinoline (25 mg, 0.15 mmol, 1 eq), chloroacetyl chloride (36 μL, 0.45 mmol, 3 eq), pyridine (36 μL, 0.45 mmol, 3 eq), anhydrous CH_2_Cl_2_ (1.5 mL, 0.10 M amine). The crude residue was purified with flash chromatography (eluting with 0-20% EtOAc in hexanes) give **compound 3** as a white solid (9.6 mg, 27%).

^1^H NMR (600 MHz, CDCl3) δ 7.10 (s, 1H), 6.78–6.77 (m, 2H), 4.83 (s, 1H), 4.33–3.99 (m, 3H), 3.82 (s, 3H), 2.56–2.38 (m, 3H), 1.14 (d, J = 5.9 Hz, 3H). LRMS–ESI+ (*m/z*): [M+H]^+^ expected for C_13_H_16_ClNO_2_ 254.1 *m/z*; found, 254.3.

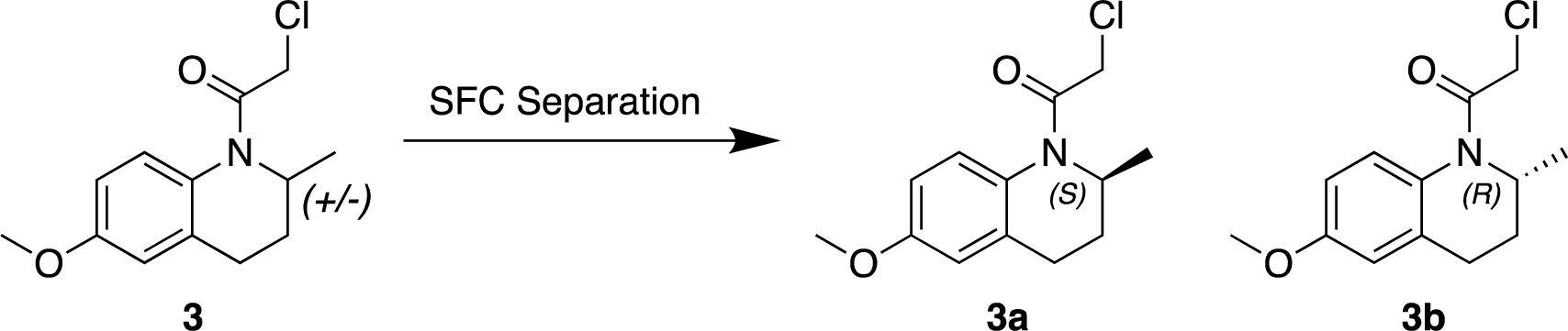

### 2-chloro-1-[(2S)-6-methoxy-2-methyl-3,4-dihydro-2H-quinolin-1-yl]ethanone (3a) 2-chloro-1-[(2*R*)-6-methoxy-2-methyl-3,4-dihydro-2*H*-quinolin-1-yl]ethenone (3b)

The racemate, compound **3** (0.0700 g, 0.273 mmol), was separated by chiral SFC separation. Column: Regis (S,S) Whelk-O 1 (Specifications: 250mm*25mm,10μm); mobile phase: CO_2_/Isopropyl alcohol (A/B); 35%-65% B, 63-minute run). **Compounds 3a** (29.5 mg, 83.4% yield) and **3b** (28.0 mg, 48.4% yield) appeared as light yellow oils. **3b** eluted off the column first (retention time: 1.51 minutes) followed by **3a** (retention time: 1.96 minutes). Vibrational circular dichroism (VCD) analysis was performed to determine absolute stereochemistry. Compound **3a**: ^1^H NMR: (CDCl_3,_ 400 MHz) *δ* 7.10 (s, 1H), 6.79 - 6.76 (m, 2H), 4.83 (s, 1H), 4.17 (d, *J* = 12.4 Hz,1H), 4.06 (d, *J* = 12.4 Hz,1H), 3.83 (s, 3H), 2.59 - 2.36 (m, 3H), 1.31 - 1.27 (m, 1H), 1.14 (d, *J* = 6.4 Hz, 3H). LRMS–ESI+ (*m/z*): [M+H]^+^ expected for C_13_H_16_ClNO_2_ 254.1 *m/z*; found 254.1. Compound **3b**: ^1^H NMR: (CDCl_3_, 400 MHz) *δ* 7.10 (s, 1H), 6.79–6.75 (m, 2H), 4.82 (s, 1H), 4.17 (d, *J* = 12.4 Hz,1H), 4.06 (d, *J* = 12.0 Hz,1H), 3.83 (s, 3H), 2.59 - 2.36 (m, 3H), 1.34 1.23 (m, 1H), 1.14 (d, *J* = 6.4 Hz, 3H). LRMS–ESI+ (*m/z*): [M+H^+^] expected for C_13_H_16_ClNO_2_ 254.1 *m/z*; found 254.1.

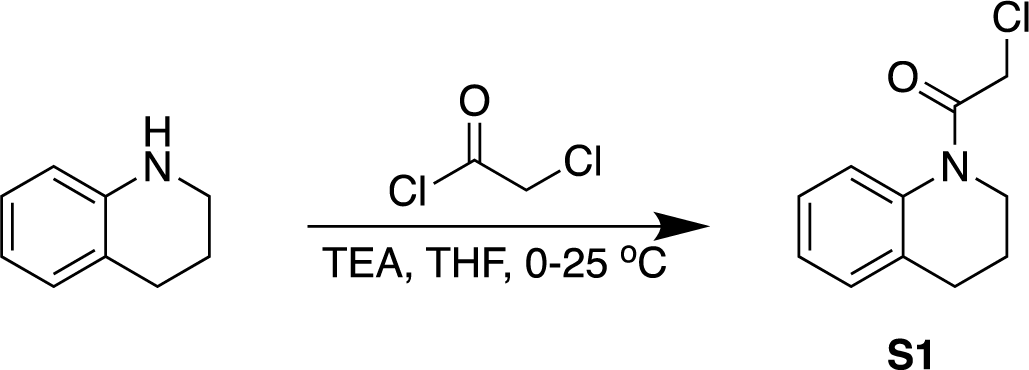

### 2-chloro-1-(3,4-dihydro-2H-quinolin-1-yl)ethanone (S1)

General procedure A was used with the following reagents: 1,2,3,4-tetrahydroquinoline (0.200 g, 1.50 mmol, 1.0 eq), tetrahydrofuran (10 mL), TEA (0.460 g, 4.50 mmol, 3.0 eq) and 2-chloroacetyl chloride (0.200 g, 1.80 mmol, 1.2 eq). The reaction was stirred at 25 °C for 1.5 hours. The crude residue was purified by prep-HPLC (Mobile phase: 10 mM NH_4_HCO_3_/MeCN (A/B); 33%-63% B, 10-minute run) to afford **compound S1** (81.0 mg, 82.0% yield) as a white solid. ^1^H NMR (CDCl_3_, 400 MHz) *δ* 7.24 - 7.18 (m, 4H), 4.24 (s, 2H), 3.84 (t, *J* = 6.8 Hz, 2H), 2.75 (t, *J* = 7.2 Hz, 2H), 2.03 - 2.00 (m, 2H). LRMS–ESI+ (*m/z*): [M+H^+^] expected for C_11_H_12_ClNO 210.1 *m/z*; found 210.0.

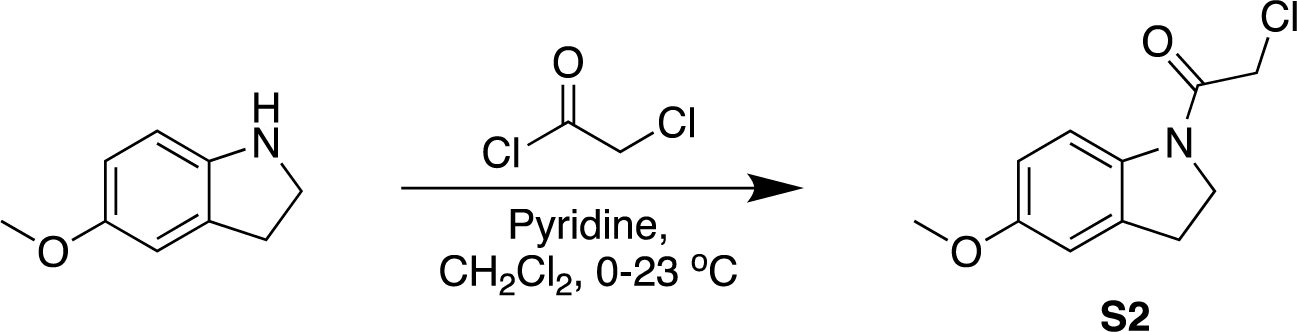

### 2-Chloro-1-(5-methoxyindolin-1-yl)ethan-1-one (S2)

General procedure A was used with the following reagents: 5-methoxyindoline (0.020 g, 0.13 mmol, 1.0 eq), chloroacetyl chloride (31 μL, 0.39 mmol, 3.0 eq), pyridine (32 μL, 0.39 mmol, 3.0 eq), CH_2_Cl_2_ (2.6 mL). The crude residue was purified with flash chromatography (eluting with 0-5% EtOAc in hexanes) to give **compound S2** as a yellow solid (10.7 mg, 37%). Spectra matched those previously reported^9^.

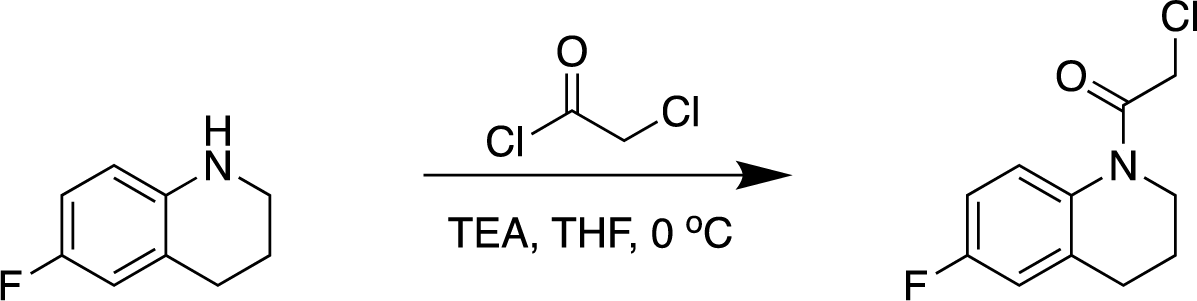

### 2-chloro-1-(6-fluoro-3,4-dihydroquinolin-1(2H)-yl)ethanone (S3)

General procedure B was used with the following reagents: 6-fluoro-1,2,3,4-tetrahydroquinoline (0.300 g, 1.98 mmol, 1.0 eq), THF (1 mL), TEA (0.402 g, 3.96 mmol, 2.0 eq) and 2-chloroacetyl chloride (0.224 g, 1.98 mmol, 1.0 eq). The mixture was stirred at 0 °C for 10 minutes. The crude product was purified by reversed-phase flash chromatography (MeCN in H_2_O, 0.1% formic acid) to afford **compound S3** (120 mg, 26.3% yield) as a brown solid. ^1^H NMR: (CDCl_3_, 400 MHz) δ 7.26 - 7.20 (m, 1H), 5.96 - 6.91 (m, 2H), 4.20 (s, 2H), 3.82 (t, *J* = 6.4 Hz, 2H), 2.77 – 2.73 (m, 2H), 2.13 - 1.90 (m, 2H). LRMS–ESI+ (*m/z*): [M+H^+^] expected for C_11_H_11_ClFNO 228.1 *m/z*; found 227.8.

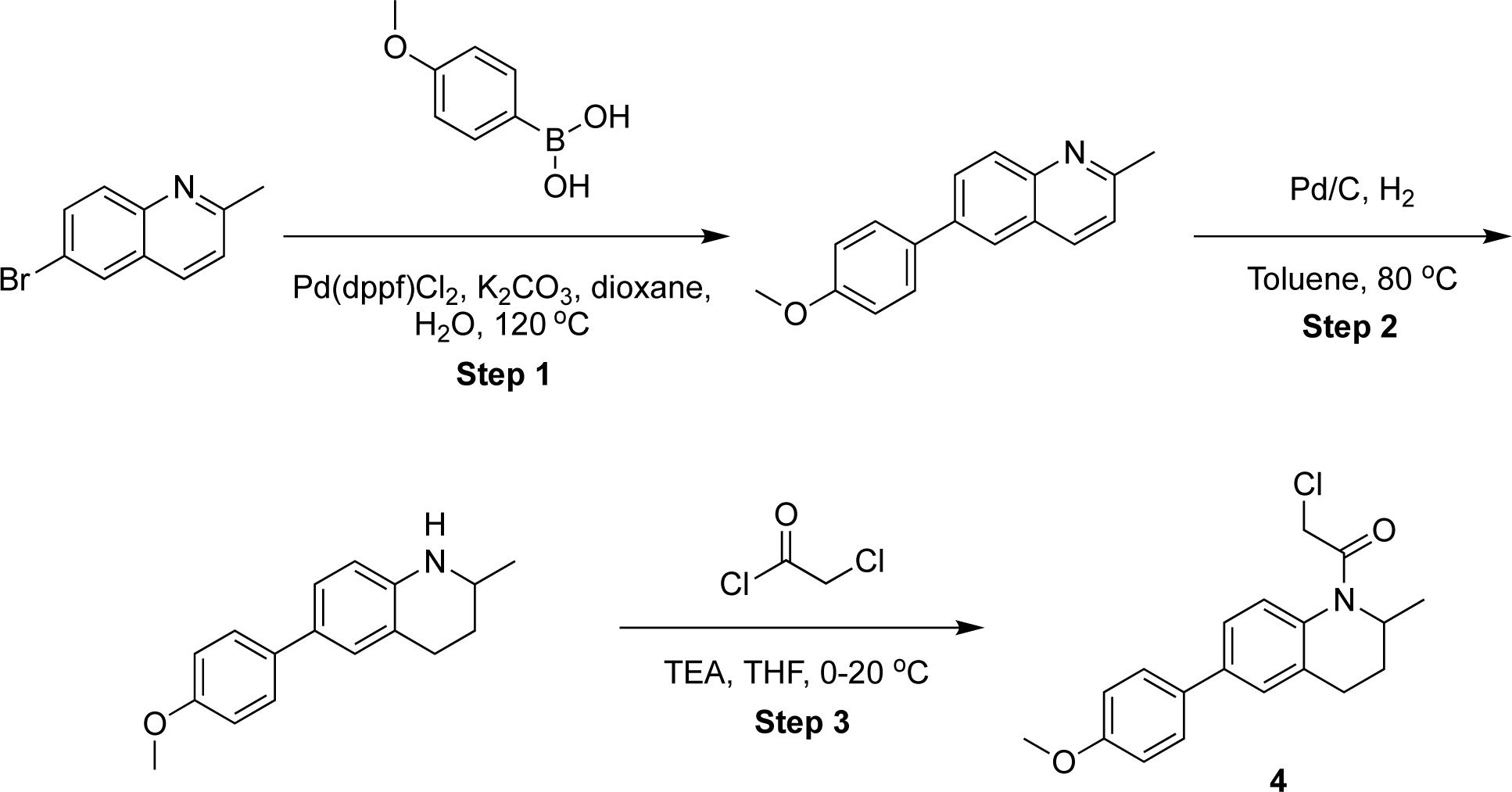

### 2-chloro-1-[6-(4-methoxyphenyl)-2-methyl-3,4-dihydro-2H-quinolin-1-yl]ethenone (4)

**Step 1.** A mixture of 6-bromo-2-methyl-quinoline (5 g, 22.51 mmol, 1 eq), 4-methoxyphenyl boronic acid (4.11 g, 27.02 mmol, 1.2 eq), Pd(dppf)Cl_2_ (1.65 g, 2.25 mmol, 0.1 eq), and K_2_CO_3_ (6.22 g, 45.03 mmol, 2 eq) in dioxane (80 mL) and H_2_O (20 mL) was degassed and purged with N_2_ 3 times. The mixture was stirred at 120 °C for 12 hours under an N_2_ atmosphere. The mixture was then poured into H_2_O (100 mL) and extracted with ethyl acetate (100 mL x 3). The combined organic layers were washed with brine (150 mL), dried over Na_2_SO_4_, filtered and concentrated under reduced pressure. The residue was purified by flash chromatography (eluting with 0-50% EtOAc in petroleum ether) to yield 6-(4-methoxyphenyl)-2-methyl-quinoline (5.5 g, 22.06 mmol, 97.99% yield) as a yellow solid. LRMS–ESI+ (*m/z*): [M+H^+^] expected for C_17_H_15_NO 250.1 *m/z*; found 250.1.

**Step 2.** General procedure A was used with the following reagents: 6-(4-methoxyphenyl)-2-methyl-quinoline (1 g, 4.01 mmol, 1 eq), toluene (10 mL), and Pd/C (213.43 mg, 10%). The mixture was stirred under H_2_ (50 Psi) at 80 °C for 24 hours, then filtered and concentrated to yield 6-(4-methoxyphenyl)-2-methyl-1,2,3,4-tetrahydroquinoline (1 g, crude) as a white solid. LRMS–ESI+ (*m/z*): [M+H^+^] expected for C_17_H_19_NO 254.2 *m/z*; found 254.3.

**Step 3.** General procedure B was used with the following reagents: 6-(4-methoxyphenyl)-2-methyl-quinoline (300 mg, 1.20 mmol, 1 eq), DCM (5 mL), TEA (243.53 mg, 2.41 mmol, 334.98 μL, 2 eq) and 2-chloroacetyl chloride (176.68 mg, 1.56 mmol, 124.60 μL, 1.3 eq). The mixture was stirred at 20 °C for 1 hour. Upon completion, the reaction mixture was filtered and the filtrate was concentrated. The residue was purified by prep-HPLC (Mobile phase: 10 mM NH_4_HCO_3_/MeCN (A/B); 35%-65% B, 8-minute run) to yield **compound 4** (83.9 mg, 21.1% yield) as a yellow solid. ^1^H NMR: (400 MHz, methanol-d4)δ 7.59-7.54 (m, 2H), 7.47 (s, 2H), 7.42-7.29 (m, 1H), 7.00 (d, *J* = 8.6 Hz, 2H), 4.8-4.68 (m, 1H), 4.40-4.32 (m, 1H), 4.28-4.19 (m, 1H), 3.83 (s, 3H), 2.80-2.68 (m, 1H), 2.63-2.40 (m, 2H), 1.49-1.25 (m, 1H), 1.16 (d, *J* = 6.4 Hz, 3H). LRMS–ESI+ (*m/z*): [M+H^+^] expected for C_19_H_20_ClNO_2_ 330.1 *m/z*; found 330.0.

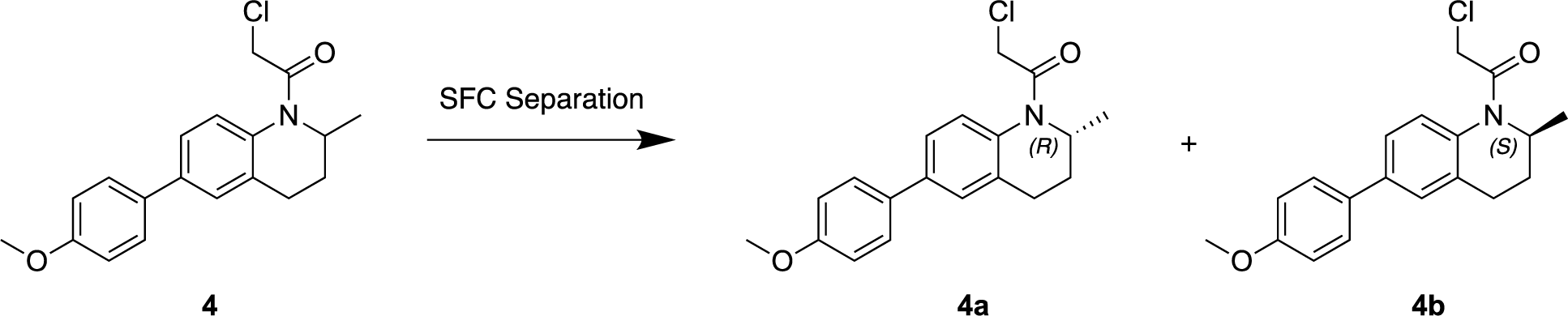

### 2-chloro-1-[(2R)-6-(4-methoxyphenyl)-2-methyl-3,4-dihydro-2H-quinolin-1-yl]ethenone (3a) and 2-chloro-1-[(2S)-6-(4-methoxyphenyl)-2-methyl-3,4-dihydro-2H-quinolin-1-yl]ethenone (3b)

The racemate, compound **3** (0.0700 g, 0.273 mmol), was separated by chiral SFC separation. Column: Regis (S,S) Whelk-O 1 (Specifications: 250mm*25mm,10μm); mobile phase: CO_2_/Methanol (A/B), 40% B) to yield compounds **4a** (31.4 mg, 7.76% yield) and **4b** (53.50 mg, 13.42% yield) as white solids. **4b** eluted off the column first (retention time 2.15 min) followed by **4b** (retention time 2.72 min). Vibrational circular dichroism (VCD) was used to determine absolute stereochemistry. Compound **4a**: ^1^H NMR: (400 MHz, methanol-d_4_) δ 7.56 (dd, *J* = 2.6, 8.9 Hz, 2H), 7.47 (br s, 2H), 7.36 (br s, 1H), 6.99 (dd, *J* = 2.6, 8.9 Hz, 2H), 4.74 (br d, *J* = 2.4 Hz, 1H), 4.40-4.30 (m, 1H), 4.29-4.19 (m, 1H), 3.83 (d, *J* = 2.8 Hz, 3H), 2.73 (br d, *J* = 12.9 Hz, 1H), 2.64-2.36 (m, 2H), 1.36 (br d, *J* = 3.3 Hz, 1H), 1.16 (dd, *J* = 2.5, 6.4 Hz, 3H). LRMS–ESI+ (*m/z*): [M+H^+^] expected for C_19_H_20_ClNO_2_ 330.1 *m/z*; found 330.1. Compound **4b**: ^1^H NMR: (400 MHz, methanol-d_4_) δ 7.59-7.54 (m, 2H), 7.50-7.45 (m, 2H), 7.36 (br s, 1H), 7.04-6.96 (m, 2H), 4.75 (br d, *J* = 5.4 Hz, 1H), 4.40-4.31 (m, 1H), 4.28-4.20 (m, 1H), 3.83 (s, 3H), 2.79-2.69 (m, 1H), 2.64-2.38 (m, 2H), 1.44-1.28 (m, 1H), 1.16 (d, *J* = 6.4 Hz, 3H). LRMS–ESI+ (*m/z*): [M+H^+^] expected for C_19_H_20_ClNO_2_ 330.1 *m/z*; found 330.1.

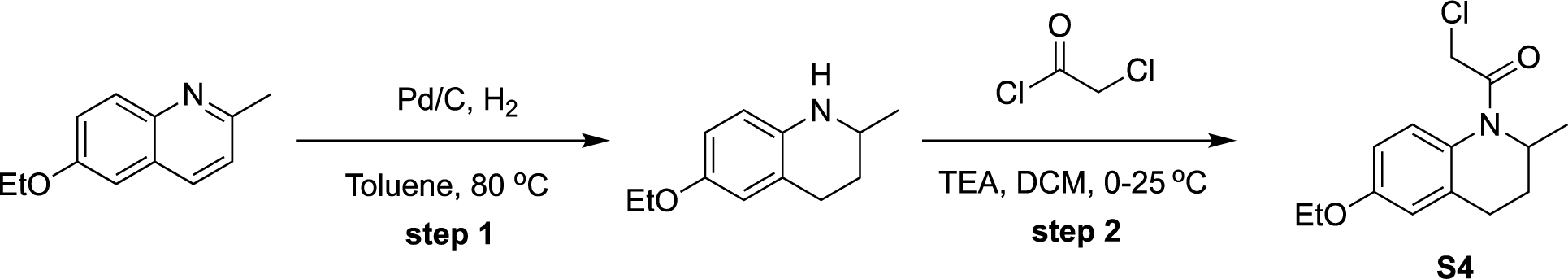

### 2-chloro-1-(6-ethoxy-2-methyl-3,4-dihydro-2H-quinolin-1-yl)ethenone (S4)

**Step 1.** General procedure A was used with the following reagents: 6-ethoxy-2-methyl-quinoline (0.5 g, 2.67 mmol, 1 eq), toluene (5 mL), and Pd/C (0.1 g, 10%). The mixture was stirred under H_2_ at 80 °C for 12 h, then filtered and concentrated to yield 6-ethoxy-2-methyl-1,2,3,4-tetrahydroquinoline (400 mg, crude).

**Step 2.** General procedure B was used with the following reagents: 6-ethoxy-2-methyl-1,2,3,4-tetrahydroquinoline (200 mg, 1.07 mmol, 1 eq), DCM (3 mL), TEA (216.17 mg, 2.14 mmol, 297.35 μL, 2 eq) and 2-chloroacetyl chloride (156.83 mg, 1.39 mmol, 110.60 μL, 1.3 eq). The mixture was stirred at 25 °C for 2 hours, then concentrated and purified by prep-HPLC (Mobile phase: 10 mM NH_4_HCO_3_/MeCN (A/B); 30%-60% B, 8-minute run) to yield **S4** (62.4 mg, 233.05 μmol, 21.82% yield, 100% purity) as a brown solid. ^1^H NMR: (400 MHz, methanol-d_4_) δ 7.18 (br s, 1H), 6.87-6.74 (m, 2H), 4.73 (br s, 1H), 4.26 (br d, *J* = 12.4 Hz, 1H), 4.15 (br d, *J* = 11.9 Hz, 1H), 4.05 (q, *J* = 7.0 Hz, 2H), 2.60 (br d, *J* = 11.1 Hz, 1H), 2.54-2.31 (m, 2H), 1.39 (t, *J* = 6.9 Hz, 3H), 1.28-1.18 (m, 1H), 1.11 (br d, *J* = 6.4 Hz, 3H). LRMS–ESI+ (*m/z*): [M+H^+^] expected for C_14_H_18_ClNO_2_ 268.1 *m/z*; found 268.0.

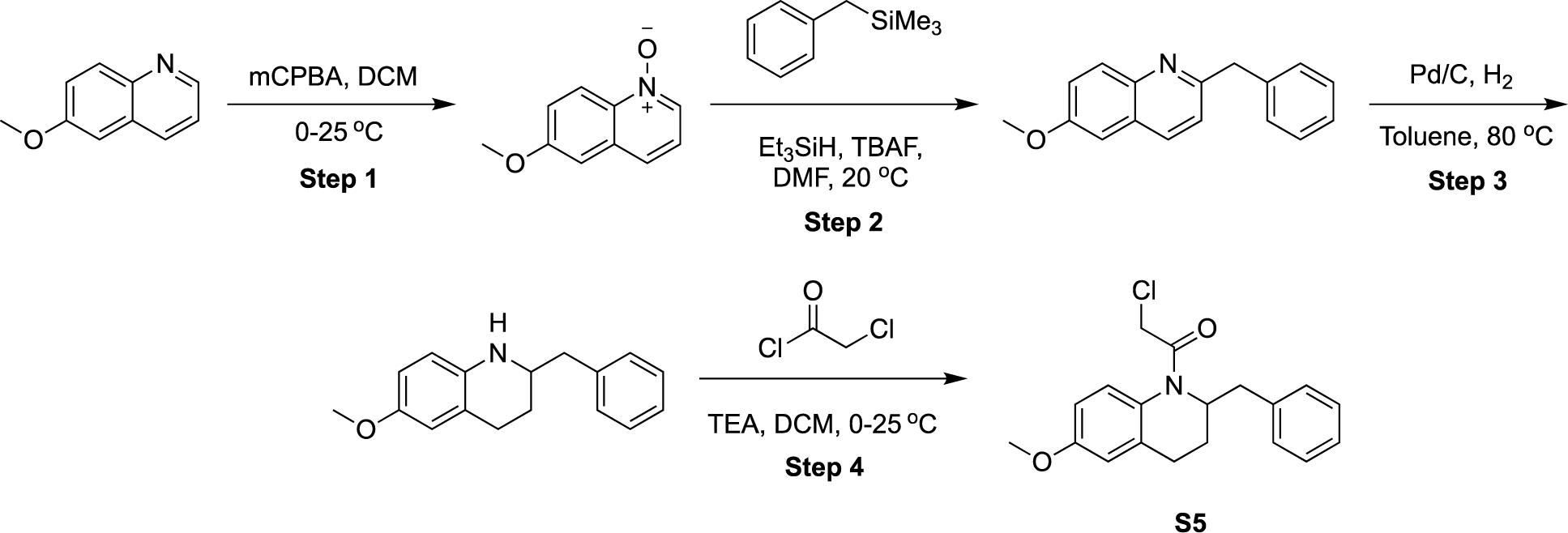

### 1-(2-benzyl-6-methoxy-3,4-dihydro-2H-quinolin-1-yl)-2-chloro-ethanone (S5)

**Step 1.** 6-methoxyquinoline (1.74 mL, 12.56 mmol, 1.0 eq) was dissolved in DCM (20 mL) and m-CPBA (3.25 g, 15.08 mmol, 80% purity, 1.2 eq) was added at 0 °C. The temperature was allowed to rise to 25 °C and the mixture was stirred for 12 h. Then, the pH was adjusted to 10 with KOH, and the mixture was stirred for an additional 30 minutes at 25 °C. The reaction was then diluted with H_2_O (20 mL) and extracted with DCM (10 mL x 3). The organic phase was separated, washed with brine (20 mL), dried over Na_2_SO_4_, filtered and concentrated under reduced pressure to yield 6-methoxy-1-oxido-quinolin-1-ium (2.1 g, crude) as a colorless oil.

**Step 2.** 6-methoxy-1-oxido-quinolin-1-ium (1 g, 5.71 mmol, 1.0 eq), benzyltrimethylsilane (2.81 g, 17.12 mmol, 3.0 eq), TBAF (1 M, 1.14 mL, 0.2 eq), and Et_3_SiH (3.32 g, 28.54 mmol, 4.56 mL, 5.0 eq) were dissolved in DMF (10 mL). The mixture was degassed and purged with N_2_ 3 times, and then stirred at 20 °C for 3 hours under N_2_. The reaction mixture was then diluted with H_2_O (10 mL) and extracted with ethyl acetate (10 mL x 3). The organic phase was separated, washed with brine (20 mL), dried over Na_2_SO_4_, filtered and concentrated under reduced pressure. The residue was purified by flash chromatography (eluting with 0-50% EtOAc in petroleum ether) to yield 2-benzyl-6-methoxy-quinoline (1 g, 4.01 mmol, 70.27% yield) as a yellow solid.

**Step 3.** General procedure A was used with the following reagents: 2-benzyl-6-methoxy-quinoline (500 mg, 2.01 mmol, 1 eq), toluene (5 mL) and added Pd/C (50 mg, 10%). The mixture was stirred under H_2_ at 80 °C for 24 hours, filtered and then concentrated to yield 2-benzyl-6-methoxy-1,2,3,4-tetrahydroquinoline (300 mg, crude) as a white solid.

**Step 4.** General procedure B was used with the following reagents: 2-benzyl-6-methoxy-1,2,3,4-tetrahydroquinoline (300 mg, 1.18 mmol, 1.0 eq), DCM (3 mL), TEA (329.65 μL, 2.37 mmol, 2.0 eq) and 2-chloroacetyl chloride (103.75 μL, 1.30 mmol, 1.1 eq). The mixture was stirred at 25 °C for 2 hours, concentrated then purified by prep-HPLC (Mobile phase: 10 mM NH_4_HCO_3_/MeCN (A/B); 45%-75% B, 8-minute run) to yield **S5** (21.3 mg, 64.21 μmol, 5.42% yield, 99.43% purity) as a white solid. ^1^H NMR: (400 MHz, methanol-d_4_) δ 7.28-7.21 (m, 2H), 7.20-7.08 (m, 4H), 6.88-6.76 (m, 2H), 5.04-4.91 (m, 1H), 4.31-4.19 (m, 1H), 4.18-4.05 (m, 1H), 3.81 (s, 3H), 2.88 (br dd, *J* = 5.3, 12.4 Hz, 1H), 2.72-2.39 (m, 3H), 2.26-2.13 (m, 1H), 1.49-1.35 (m, 1H). LRMS–ESI+ (*m/z*): [M+H^+^] expected for C_19_H_20_ClNO_2_ 330.1 *m/z*; found 330.1.

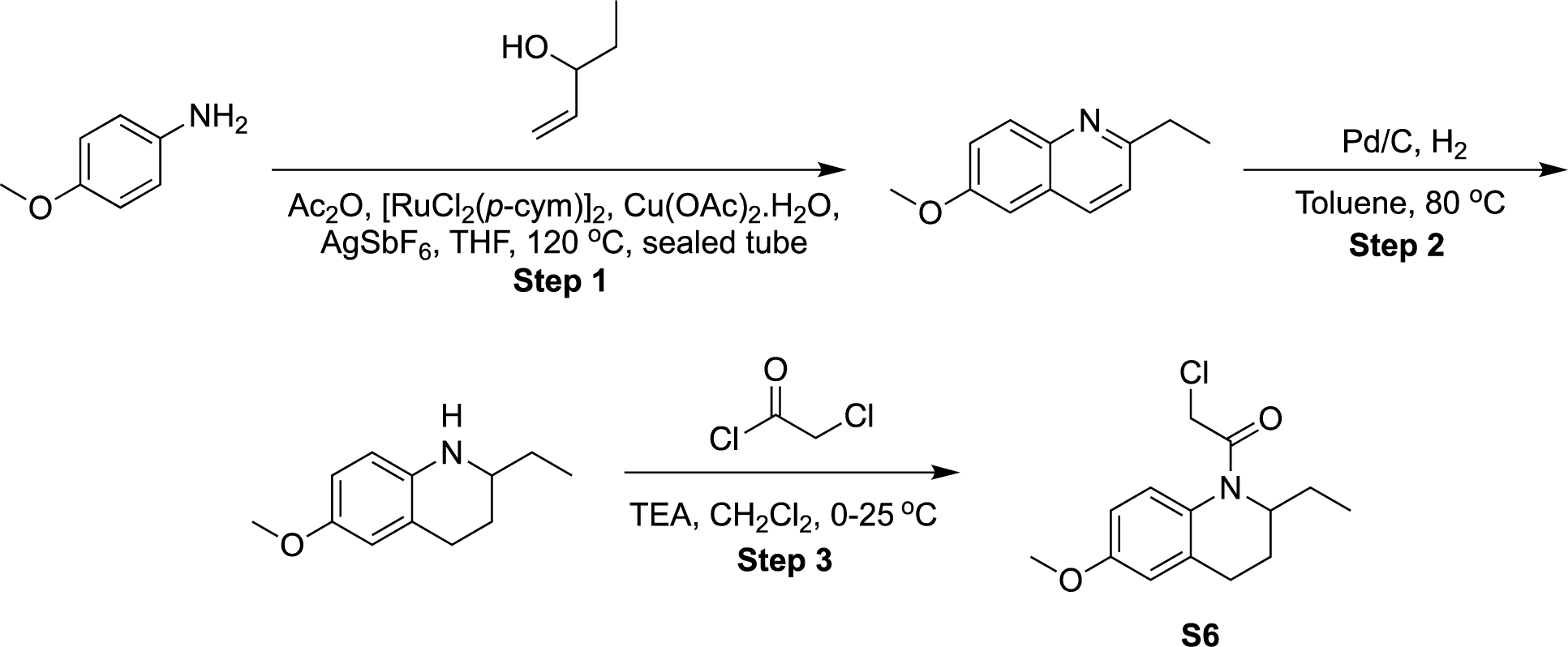

### 2-chloro-1-(2-ethyl-6-methoxy-3,4-dihydro-2H-quinolin-1-yl)ethenone (S6)

**Step 1.** A mixture of 4-methoxyaniline (500 mg, 4.06 mmol, 1.0 eq), pent-1-en-3-ol (699.39 mg, 8.12 mmol, 834.60 μL, 2.0 eq), Ac_2_O (762.62 μL, 8.12 mmol, 2.0 eq), Cu(OAc)_2_.H_2_O (1.62 g, 8.12 mmol, 2.0 eq), AgSbF_6_ (697.55 mg, 2.03 mmol, 0.5 eq), and [RuCl_2_(*p*-cym)]_2_ (124.32 mg, 203.00 μmol, 0.05 eq) in THF (5 mL) was degassed and purged with N_2_ 3 times. The mixture was then stirred in a sealed tube at 120 °C for 24 horus under an N_2_ atmosphere. The reaction mixture was filtered, and the filtrate was poured into H_2_O (10 mL), followed by extraction with CH_2_Cl_2_ (10 mL x 3). The combined organic layers were washed with brine (20 mL), dried over Na_2_SO_4_, filtered and concentrated under reduced pressure. The residue was purified by flash chromatography (eluting with 50-100% EtOAc in petroleum ether) to yield 2-ethyl-6-methoxy-quinoline (200 mg, 26.31% yield) as a brown oil.

**Step 2.** General procedure A was used with the following reagents: 2-ethyl-6-methoxy-quinoline (200 mg, 1.07 mmol, 1 eq), toluene (2 mL) and Pd/C (20 mg, 10%). The mixture was stirred under H_2_ at 80 °C for 24 hours, then filtered and concentrated to yield 2-ethyl-6-methoxy-1,2,3,4-tetrahydroquinoline (190 mg, crude) as colorless oil. LRMS–ESI+ (*m/z*): [M+H^+^] expected for C_12_H_17_NO 192.1 *m/z*; found 192.0.

**Step 3.** General procedure B was used with the following reagents: 2-ethyl-6-methoxy-1,2,3,4-tetrahydroquinoline (190 mg, 993.37 μmol, 1.0), DCM (3 mL), TEA (276.53 μL, 1.99 mmol, 2.0 eq) and 2-chloroacetyl chloride (87.03 μL, 1.09 mmol, 1.1 eq). The mixture was stirred at 25 °C for 2 hours, concentrated under reduced pressure, and purified by prep-HPLC (Mobile phase: 10 mM NH_4_HCO_3_/MeCN (A/B); 45%-75% B, 8-minute run) to yield **S6** (66.5 mg, 23.52% yield) as a white solid. ^1^H NMR: (400 MHz, methanol-d_4_) δ 7.21 (br d, *J* = 8.1 Hz, 1H), 6.88-6.78 (m, 2H), 4.76-4.59 (m, 1H), 4.34-4.07 (m, 2H), 3.81 (s, 3H), 2.71-2.31 (m, 3H), 1.51-1.25 (m, 3H), 0.88 (t, *J* = 7.4 Hz, 3H). LRMS–ESI+ (*m/z*): [M+H^+^] expected for C_14_H_18_ClNO_2_ 268.1 *m/z*; found 268.1.

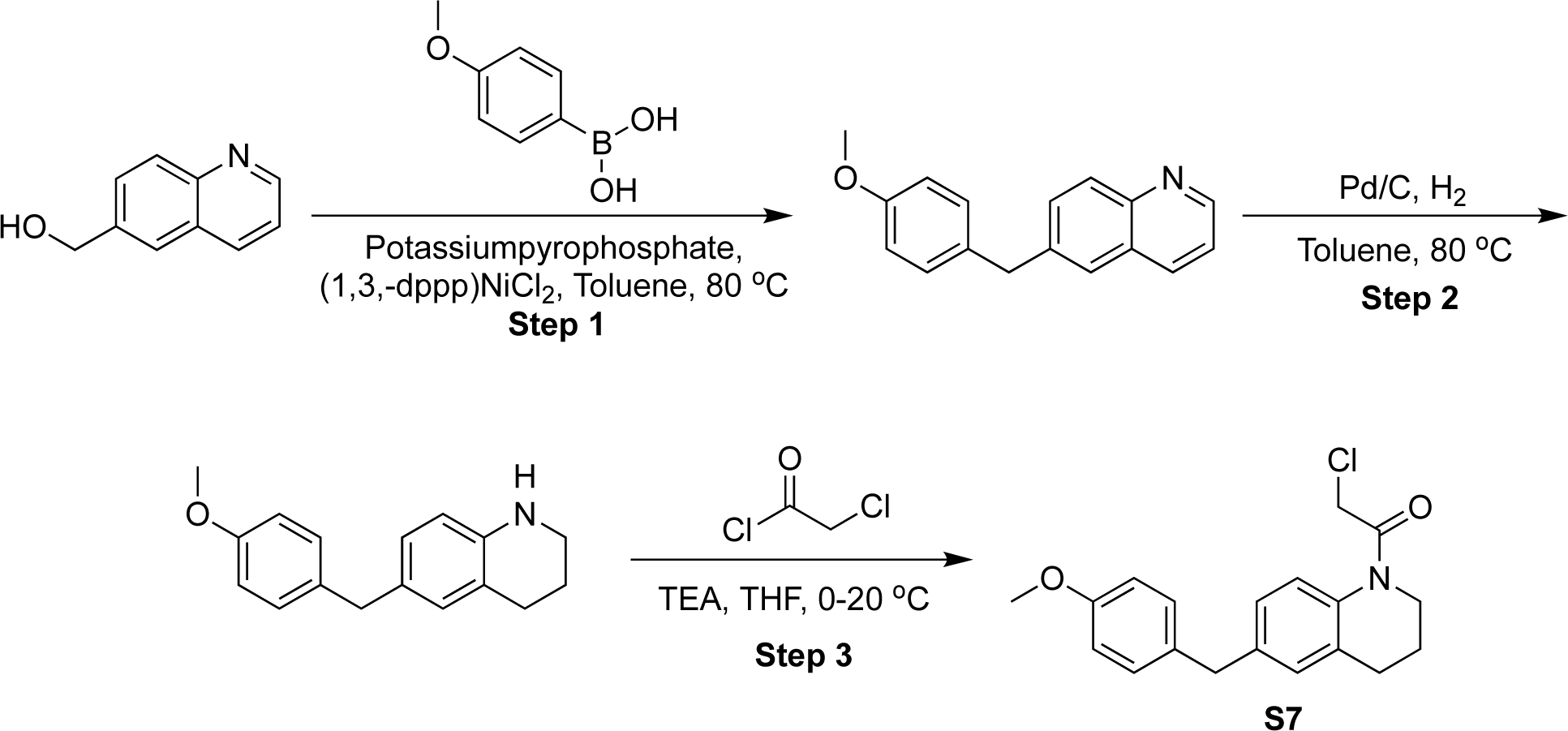

### 2-chloro-1-[6-[(4-methoxyphenyl)methyl]-3,4-dihydro-2H-quinolin-1-yl]ethanone (S7)

Step 1. 6-quinolylmethanol (500 mg, 3.14 mmol, 1.0 eq) and (4-methoxyphenyl)boronic acid (1.43 g, 9.42 mmol, 3.0 eq) were dissolved in toluene (20 mL). [1,3-Bis(diphenylphosphino)propane]dichloronickel(II) (170.26 mg, 314.10 μmol, 0.1 eq) and potassiumpyrophosphate (3.62 g, 9.42 mmol, 3.0 eq) were added and he mixture was stirred at 80 °C for 12 h. The reaction was concentrated under reduced pressure and diluted with H_2_O (10 mL). The mixture was extracted with ethyl acetate (15 mL x 3) and the combined organic layers were washed with brine (20 mL), dried over Na_2_SO_4_, filtered and concentrated under reduced pressure. The residue was purified by column chromatography (eluting with 0-25% EtOAc in petroleum ether) to yield 6-[(4-methoxyphenyl)methyl]quinoline (550 mg, 49.17% yield) as a yellow oil. LRMS–ESI+ (*m/z*): [M+H^+^] expected for C_17_H_15_NO 250.1 *m/z*; found 250.2.

**Step 2**. General procedure A was used with the following reagents: 6-[(4-methoxyphenyl)methyl]quinoline (500 mg, 2.01 mmol, 1.0 eq), toluene (5 mL), Pd/C (213.43 mg, 10%). The reaction was stirred at 80 °C for 12 hours under H_2_, then filtered and concentrated to yield 6-[(4-methoxyphenyl)methyl]-1,2,3,4-tetrahydroquinoline (480 mg, crude) as colorless oil. LRMS–ESI+ (*m/z*): [M+H^+^] expected for C_17_H_19_NO 254.2 *m/z*; found 254.2.

**Step 3**. General procedure B was used with the following reagents: 6-[(4-methoxyphenyl)methyl]-1,2,3,4-tetrahydroquinoline (250 mg, 986.82 μmol, 1.0 eq), DCM (5 mL), TEA (274.71 μL, 1.97 mmol, 2.0 eq), and 2-chloroacetyl chloride (102.18 μL, 1.28 mmol, 1.3 eq). The mixture was stirred at 20 °C for 1 hour, then filtered and concentrated. The residue was purified by prep-HPLC (Mobile phase: 10 mM NH_4_HCO_3_/MeCN (A/B); 40%-70% B, 8-minute run) to yield **compound S7** (76.8 mg, 21.99% yield) as a white solid. ^1^H NMR: (400 MHz, methanol-d4) δ 7.39-7.14 (m, 1H), 7.10 (br d, *J* = 8.5 Hz, 2H), 7.04 (br s, 2H), 6.82 (d, *J* = 8.5 Hz, 2H), 4.32 (br s, 2H), 3.86 (s, 2H), 3.79-3.75 (m, 2H), 3.75 (s, 3H), 2.69 (br s, 2H), 1.99-1.91 (m, 2H). LRMS–ESI+ (*m/z*): [M+H^+^] expected for C_19_H_20_ClNO_2_ 330.1 *m/z*; found 330.1

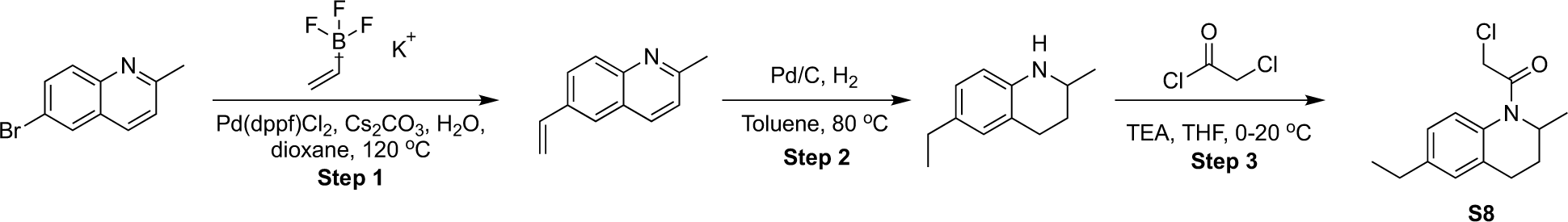

### 2-chloro-1-(6-ethyl-2-methyl-3,4-dihydro-2H-quinolin-1-yl)ethanone (S8)

**Step 1.** 6-bromo-2-methyl-quinoline (5 g, 22.51 mmol, 1.0 eq), potassium vinyltrifluoroborate (3.62 g, 27.02 mmol, 1.2 eq), Pd(dppf)Cl_2_ (1.65 g, 2.25 mmol, 0.1 eq) and Cs_2_CO_3_ (14.67 g, 45.03 mmol, 2 eq) were dissolved in dioxane (80 mL) and H_2_O (20 mL). The mixture was degassed and purged with N_2_ 3 times and then stirred at 120 °C for 12 hours under N_2_. The mixture was diluted with H_2_O (100 mL) and extracted with ethyl acetate (150 mL x 3). The combined organic layers were washed with brine (200 mL), dried over Na_2_SO_4_, filtered and concentrated under reduced pressure. The residue was purified by flash chromatography (eluting with 0-25% EtOAc in petroleum ether) to yield 2-methyl-6-vinyl-quinoline (3.3 g, 19.50 mmol, 86.62% yield) as a yellow oil. LRMS–ESI+ (*m/z*): [M+H^+^] expected for C_12_H_11_N 170.1 *m/z*; found 170.2.

**Step 2.** General procedure A was used with the following reagents: 2-methyl-6-vinyl-quinoline (500 mg, 2.95 mmol, 1.0 eq), toluene, (5 mL), and Pd/C (157.22 mg, 10%). The mixture was stirred under H_2_ at 80 °C for 12 hours, then filtered and concentrated to yield 6-ethyl-2-methyl-1,2,3,4-tetrahydroquinoline (400 mg, crude) as a yellow oil. LRMS–ESI+ (*m/z*): [M+H^+^] expected for C_12_H_17_N 176.1 *m/z*; found 176.2.

**Step 3.** General procedure B was used with the following reagents: 6-ethyl-2-methyl-1,2,3,4-tetrahydroquinoline (307.06 mg, 1.75 mmol, 1.0 eq), DCM (5 mL), TEA (487.70 μL, 3.50 mmol, 2.0 eq) and 2-chloroacetyl chloride (181.40 μL, 2.28 mmol, 1.3 eq). The mixture was stirred at 20 °C for 1 h, then filtered and concentrated under reduced pressure. The residue was purified by prep-HPLC (Mobile phase: 10 mM NH_4_HCO_3_/MeCN (A/B); 35%-65% B, 8-minute run) to yield **compound S8** (143 mg, 32.42% yield) as a yellow solid. ^1^H NMR: (400 MHz, methanol-d_4_) δ 7.20 (br s, 1H), 7.13-7.09 (m, 2H), 4.72 (br s, 1H), 4.33-4.27 (m, 1H), 4.17 (br d, *J* = 12.0 Hz, 1H), 2.65 (q, *J* = 7.6 Hz, 4H), 2.51 (br d, *J* = 10.4 Hz, 1H), 2.45-2.37 (m, 1H), 1.24 (t, *J* = 7.6 Hz, 3H), 1.12 (d, *J* = 6.4 Hz, 3H). LRMS–ESI+ (*m/z*): [M+H^+^] expected for C_14_H_18_ClNO 252.1 *m/z*; found 252.1.

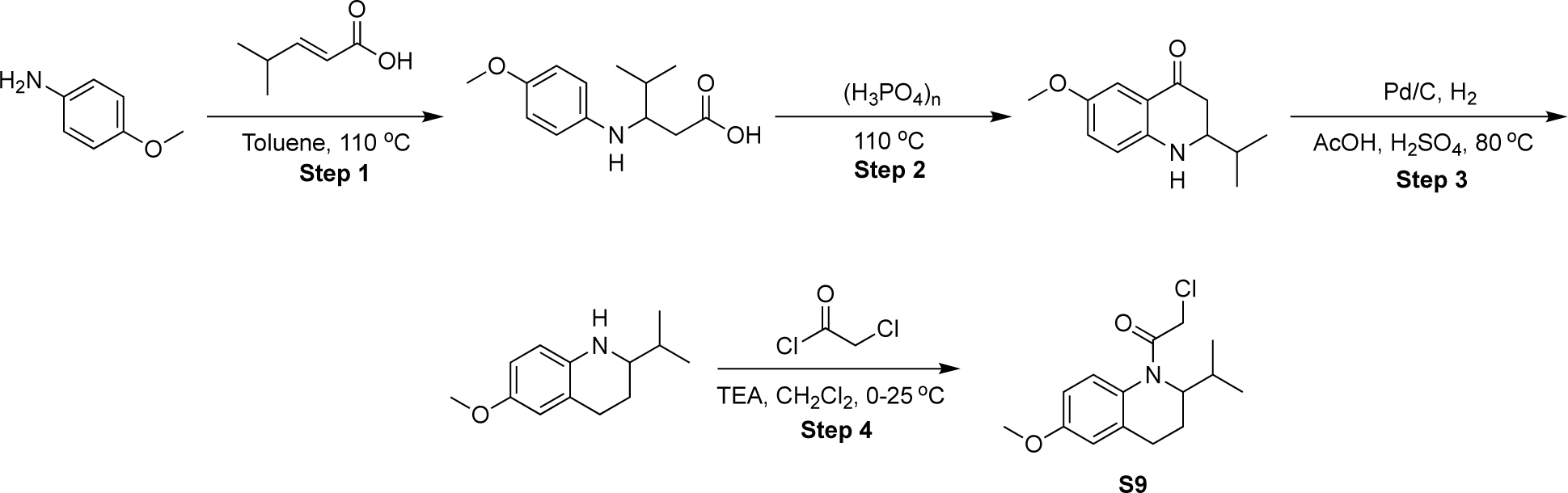

### 2-chloro-1-(2-isopropyl-6-methoxy-3,4-dihydro-2H-quinolin-1-yl)ethenone (S9)

**Step 1.** 4-methoxyaniline (1 g, 8.12 mmol, 1.0 eq) and (E)-4-methylpent-2-enoic acid (965.46 μL, 8.12 mmol, 1.0 eq) were dissolved in toluene (10 mL). The mixture was stirred at 110 °C for 12 hours. The reaction was concentrated under reduced pressure to to yield 3-(4-methoxyanilino)-4-methyl-pentanoic acid (1.9 g, crude) as a brown solid. The product was used in the next step without further purification.

**Step 2.** 3-(4-methoxyanilino)-4-methyl-pentanoic acid (1.9 g, 8.01 mmol, 1.0 eq) was suspended in polyphosphoric acid (20 mL) and then stirred at 110 °C for 6 hours. The reaction was diluted with H_2_O (10 mL) and extracted with ethyl acetate (15 mL x 3). The organic phase was separated, washed with brine (20 mL), dried over Na_2_SO_4_, filtered and concentrated. The residue was purified by flash chromatography (eluting with 0-100% EtOAc in petroleum ether) to yield 2-isopropyl-6-methoxy-2,3-dihydro-1H-quinolin-4-one (320 mg, 18.23% yield) as a yellow solid.

**Step 3.** 2-isopropyl-6-methoxy-2,3-dihydro-1H-quinolin-4-one (180 mg, 820.87 μmol, 1.0 eq) and H_2_SO_4_ (0.438 μL, 8.21 μmol, 0.01 eq) were mixed in AcOH (2 mL). Pd/C (87.36 mg, 10%) was added under an N_2_ atmosphere. The suspension was degassed and purged with H_2_ 3 times. The mixture was stirred under H_2_ (15 Psi) at 80 °C for 2 hours. The reaction mixture was then filtered and concentrated under reduced pressure. The residue was purified by prep-HPLC (Column: Phenomenex Luna C18; specifications: 100×40mm×5 um; mobile phase: H_2_O(0.04% HCl)/MeCN (A/B); 5%-30% B, 8-minute run) to yield 2-isopropyl-6-methoxy-1,2,3,4-tetrahydroquinoline (50 mg, 29.67% yield) as a white solid. LRMS–ESI+ (*m/z*): [M+H^+^] expected for C_13_H_19_NO 206.2 *m/z*; found 206.2.

**Step 4.** General procedure B was used with the following reagents: 2-isopropyl-6-methoxy-1,2,3,4-tetrahydroquinoline (15 mg, 73.07 μmol, 1.0 eq) CH_2_Cl_2_ (0.5 mL), TEA (30.51 μL, 219.20 μmol, 3.0 eq) and 2-chloroacetyl chloride (17.46 μL, 219.20 μmol, 3.0 eq). The mixture was stirred at 25 °C for 2 hours and then concentrated under reduced pressure. The residue was purified by prep-HPLC (Mobile phase: 10 mM NH_4_HCO_3_/MeCN (A/B); 37%-61% B, 8-minute run) to yield **compound S9** (4.4 mg, 21.30% yield) as a white solid. ^1^H NMR: (400 MHz, methanol-d_4_) δ 7.24 (br d, *J* = 8.3 Hz, 1H), 6.87-6.78 (m, 2H), 4.52 (q, *J* = 8.1 Hz, 1H), 4.26 (d, *J* = 12.8 Hz, 1H), 4.15-4.08 (m, 1H), 3.81 (s, 3H), 2.65 (td, *J* = 4.5, 14.7 Hz, 1H), 2.54-2.42 (m, 1H), 2.32-2.21 (m, 1H), 1.67-1.39 (m, 2H), 0.93-0.82 (m, 6H). LRMS–ESI+ (*m/z*): [M+H^+^] expected for C_15_H_20_ClNO_2_ 282.1 *m/z*; found 282.0.

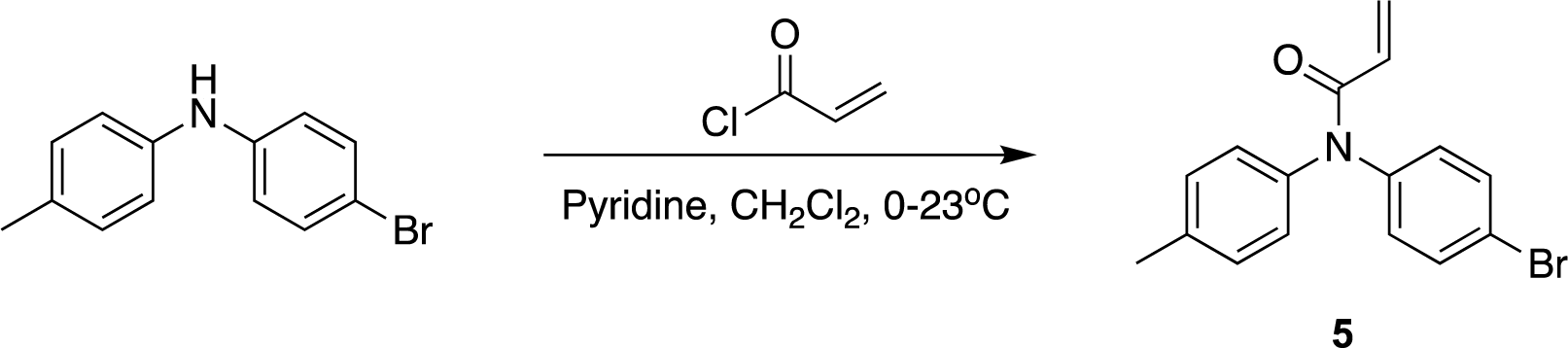

### *N*-(4-Bromophenyl)-*N*-(*p*-tolyl)acrylamide (Compound 5)

General procedure C was used with the following reagents: 4-bromo-*N*-(p-tolyl)aniline (0.020 g, 0.076 mmol, 1.0 eq), acryloyl chloride (23 μL, 0.23 mmol, 3.0 eq), pyridine (18 μL, 0.23 mmol, 3.0 eq), CH_2_Cl_2_ (0.76 mL). The crude residue was purified with flash chromatography (eluting with 0-10% EtOAc in hexanes) to give compound **5** as a yellow solid (3.3 mg, 13%).^1^H NMR (600 MHz, CDCl_3_) δ 7.46 (d, *J* = 7.8 Hz, 2H), 7.22–7.06 (m, 6H), 6.47 (dd, *J* = 16.8, 1.6 Hz, 1H), 6.18 (dd, *J* = 16.8, 10.3 Hz, 1H), 5.64 (d, *J* = 10.4 Hz, 1H), 2.38 (s, 3H). LRMS–ESI+ (*m/z*): [M+H^+^] expected for C_16_H_14_BrNO 316.0 *m/z*; found, 316.0.

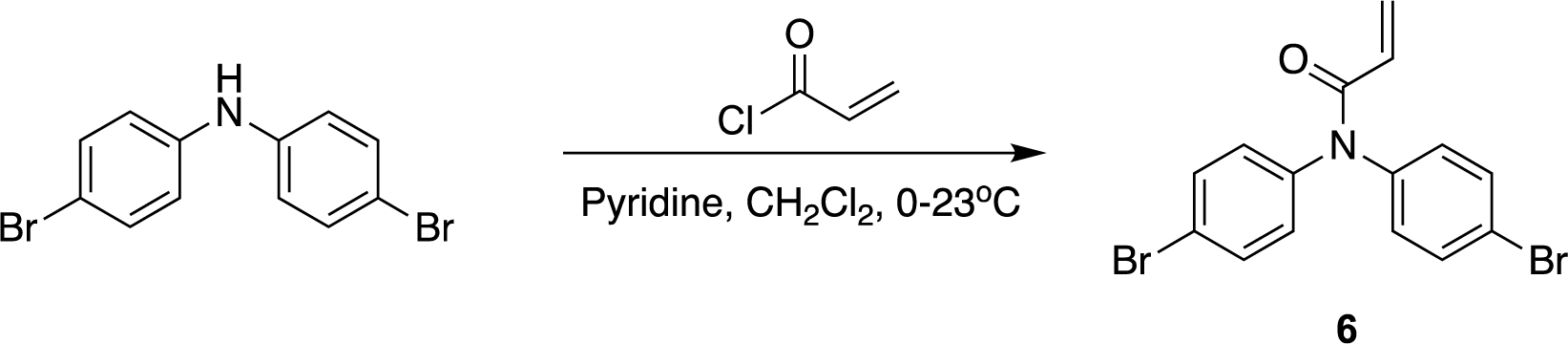

### *N*,*N*-Bis(*p*-bromophenyl)acrylamide (Compound 6)

General procedure C was used with the following reagents: bis(4-bromophenyl)amine (0.050 g, 0.15 mmol, 1.0 eq), acryloyl chloride (36 μL, 0.45 mmol, 3.0 eq), pyridine (36 μL, 0.45 mmol, 3.0 eq), anhydrous CH_2_Cl_2_ (1.5 mL, 0.10 M amine). The crude residue was purified with flash chromatography (eluting with 0-15% EtOAc in hexanes) to give compound **6** as a yellow solid (3.4 mg, 6%). ^1^H NMR (600 MHz, CDCl_3_) δ 7.51 (d, *J* = 7.7 Hz, 4H), 7.10 (d, *J* = 8.3 Hz, 4H), 6.49 (d, *J* = 16.8 Hz, 1H), 6.17 (dd, *J* = 16.7, 10.3 Hz, 1H), 5.69 (d, *J* = 10.3 Hz, 1H). LRMS–ESI+ (*m/z*): [M+H^+^] expected for C_15_H_11_Br_2_NO 379.9 *m/z*; found, 380.0.

