## Supplementary Information for "Using a function-first ‘scout fragment’-based approach to develop allosteric covalent inhibitors of conformationally dynamic helicase mechanoenzymes"

### SUPPORTING INFORMATION

TABLE OF CONTENTS

Figure S1: Site-mapping MS analysis of nsp13$^{wt}$ treated with compound **1.**
Table S1: Residues of nsp13$^{wt}$ liganded by compound **1**.
Figure S2: Biochemical characterization of nsp13 mutants.
Table S2: Kinetic parameters for nsp13 wild-type and mutant constructs. Table S1,
Figure S3: Site-mapping MS analysis of nsp13$^{C441S\ C444S}$ treated with compound **1.**
Table S3: Residues of nsp13$^{C441S\ C444S}$ liganded by compound **1**.
Figure S4: Compounds tested in first analog screen and inhibitory activity.
Figure S5: Nsp13 Covalent docking model of **3a** and **3b**.
Figure S6: Compounds tested in second analog screen and inhibitory activity.
Figure S7: Site-mapping MS analysis of nsp13$^{wt}$ treated with compound **4b.**
Table S4: Residues of nsp13$^{wt}$ liganded by compound **4b.**
Figure S8: Purification of BLM$^{636-1298}$ and WRN$^{515-1233}$ constructs.

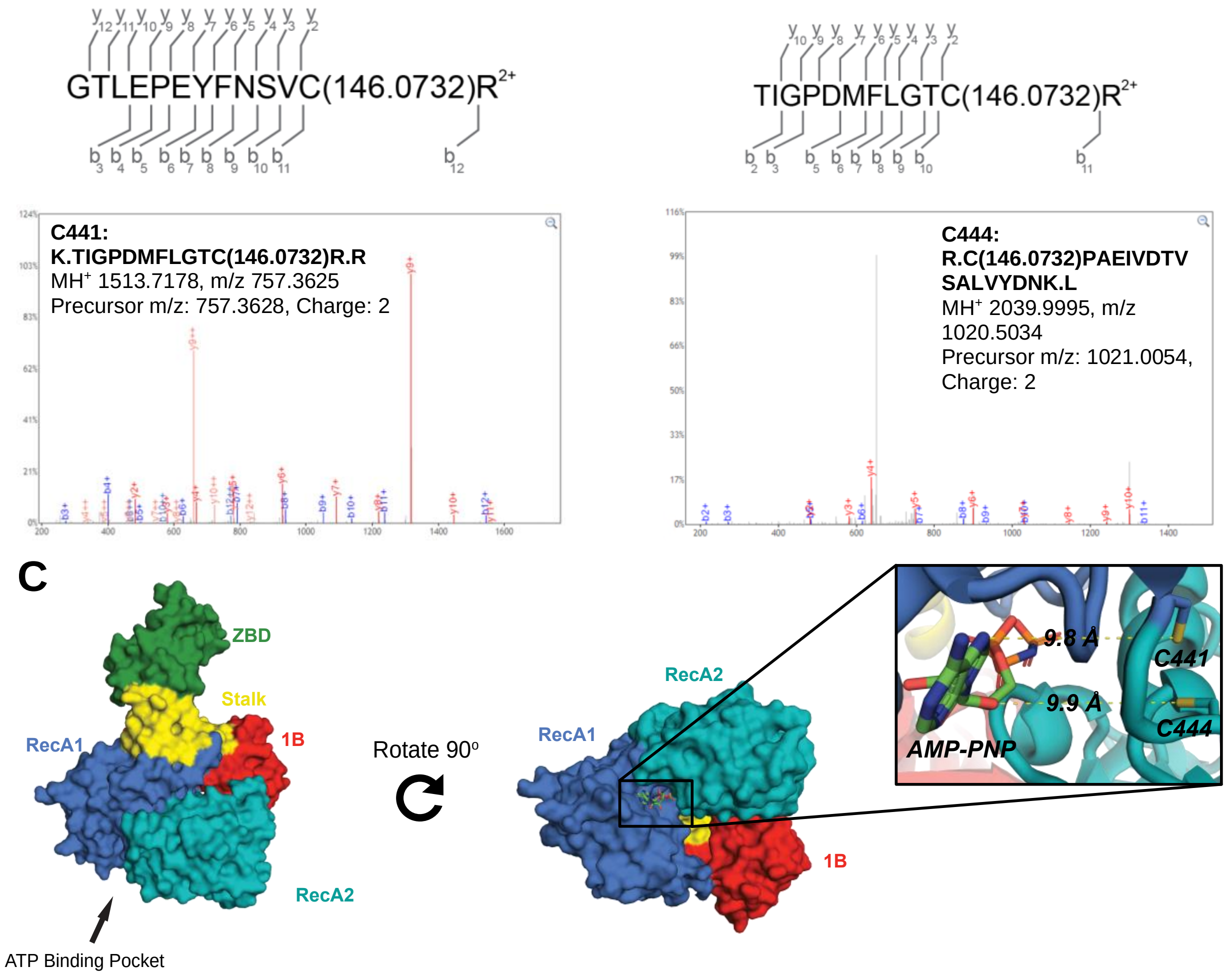


**Figure S1:** Site-mapping MS analysis of nsp13$^{wt}$ treated with compound **1** (200 μM, 24h, 4°C) a), b) Representative MS$^2$ spectra corresponding to peptides liganded by compound **1** at C441 and C444. c) Surface depiction of nsp13 with zoom on C441 and C444 residues inside the ATP binding pocket. Distance from AMP-PNP is noted (PDB: 7NN0).

| Nsp13 Cys Residue | Liganded Peptide Counts | Unliganded Peptide Counts | Total Peptide Counts | Liganding Efficiency' (%) |
|---|---|---|---|---|
| 97/112/126 | 22 | 80 | 102 | 20 |
| 309/318 | 6 | 44 | 50 | 12 |
| 342 | 3 | 3 | 6 | 50 |
| 426 | 3 | 14 | 17 | 18 |
| **441** | **11** | **5** | **16** | **69** |
| **444** | **25** | **137** | **162** | **15** |
| 556 | 4 | 96 | 100 | 4 |

**Table S1:** Site mapping MS detection of all residues of nsp13$^{wt}$ liganded by compound **1**. C441 and C444 were chosen for initial follow-up work due to location in ATP binding pocket.

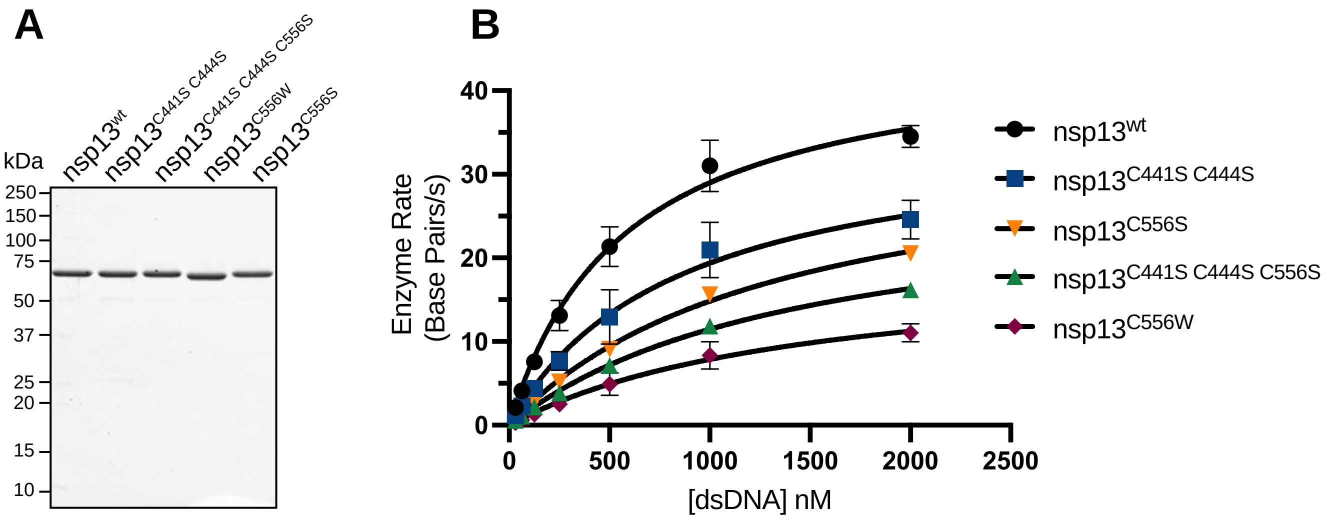


**Figure S2:** Biochemical characterization of nsp13 mutants. (a) SDS-PAGE analysis (Coomassie blue) of purified recombinant nsp13$^{wt}$ and mutants. (b) dsDNA dose-dependent helicase activity of wild-type and mutant nsp13 constructs. Rates (n=2, mean ± SD) were fit to the Michaelis Menten equation. Note: [E] = 2 nM for all constructs except nsp13$^{C556W}$, [E] = 16 nM. Values for enzyme activity parameters are shown in Table S2.

| Construct | $K_{m,\ dsDNA}$ (nM) | $k_{cat}$ $s^{-1}$ | $k_{cat}/K_m$ $s^{-1}$ $nM^{-1}$ |
|---|---|---|---|
| nsp13$^{wt}$ | 573 ± 61 | 22.8 ± 0.9 | 0.0398 |
| nsp13$^{C441S\ C444S}$ | 843 ± 156 | 17.9 ± 1.5 | 0.0212 |
| nsp13$^{C441S\ C444S\ C556S}$ | 1467 ± 84 | 14.2 ± 0.9 | 0.0097 |
| nsp13$^{C556S}$ | 1341 ± 105 | 17.4 ± 0.7 | 0.0129 |
| nsp13$^{C556W}$ | 1493 ± 321 | 1.2 ± 0.1 | 0.0008 |

**Table S2:** Kinetic parameters for nsp13 wild-type and mutant constructs. Values for half maximal rate ($K_{m,\ dsDNA}$), turnover number ($k_{cat}$) and catalytic efficiency ($k_{cat}/K_m$) were determined by fitting enzyme rates to the Michaelis Menten equation. (n=2 ± SD).

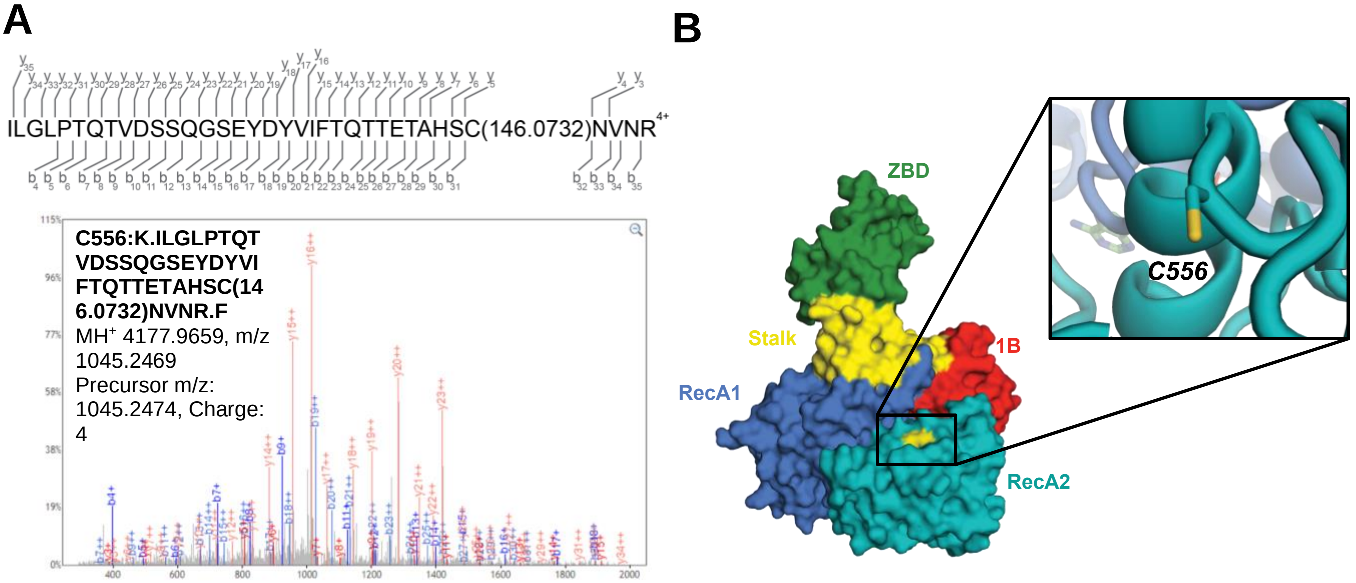


**Figure S3:** Site-mapping MS analysis of nsp13$^{C441S\ C444S}$ treated with compound **1** (200 μM, 24h, 4$^{o}$C) a), b) Representative $MS^2$ spectra corresponding to peptides liganded by compound **1** at C556. c) Surface depiction of nsp13 with zoom on C556 residue located in RecA2 domain (PDB: 7NN0).

| Nsp13 Cys Residue | Liganded Peptide Counts | Unliganded Peptide Counts | Total Peptide Counts | Liganding Efficiency' (%) |
|---|---|---|---|---|
| 26/27 | 1 | 4 | 5 | 20 |
| 30/50/55 | 3 | 7 | 10 | 30 |
| 84 | 3 | 16 | 19 | 15 |
| 309/318 | 2 | 104 | 106 | 2 |
| 330 | 2 | 1 | 3 | 67 |
| 342 | 2 | 3 | 5 | 40 |
| 426 | 7 | 24 | 31 | 23 |
| **556** | **64** | **11** | **75** | **85** |
| 574 | 3 | 54 | 57 | 5 |

**Table S3:** Residues of nsp13$^{C441S\ C444S}$ liganded by compound **1**.

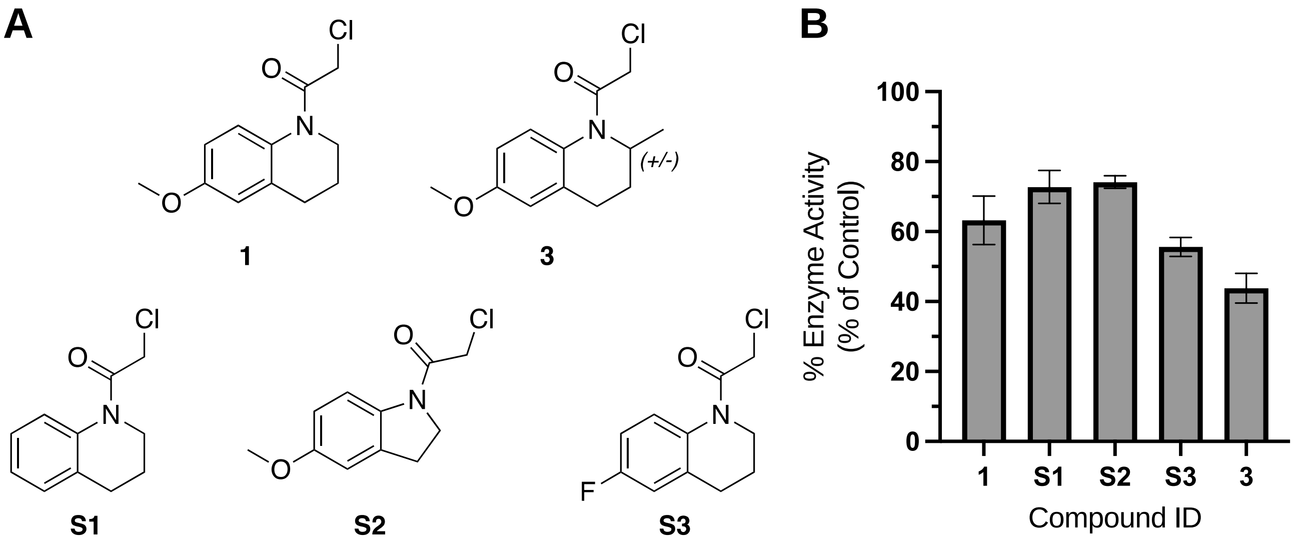


**Figure S4:** Compounds tested in first analog screen and inhibitory activity (a) Chemical structures of compounds tested. (b) Testing of compound **1** analogs (200 µM, 4h incubation, 4$^{o}$C)

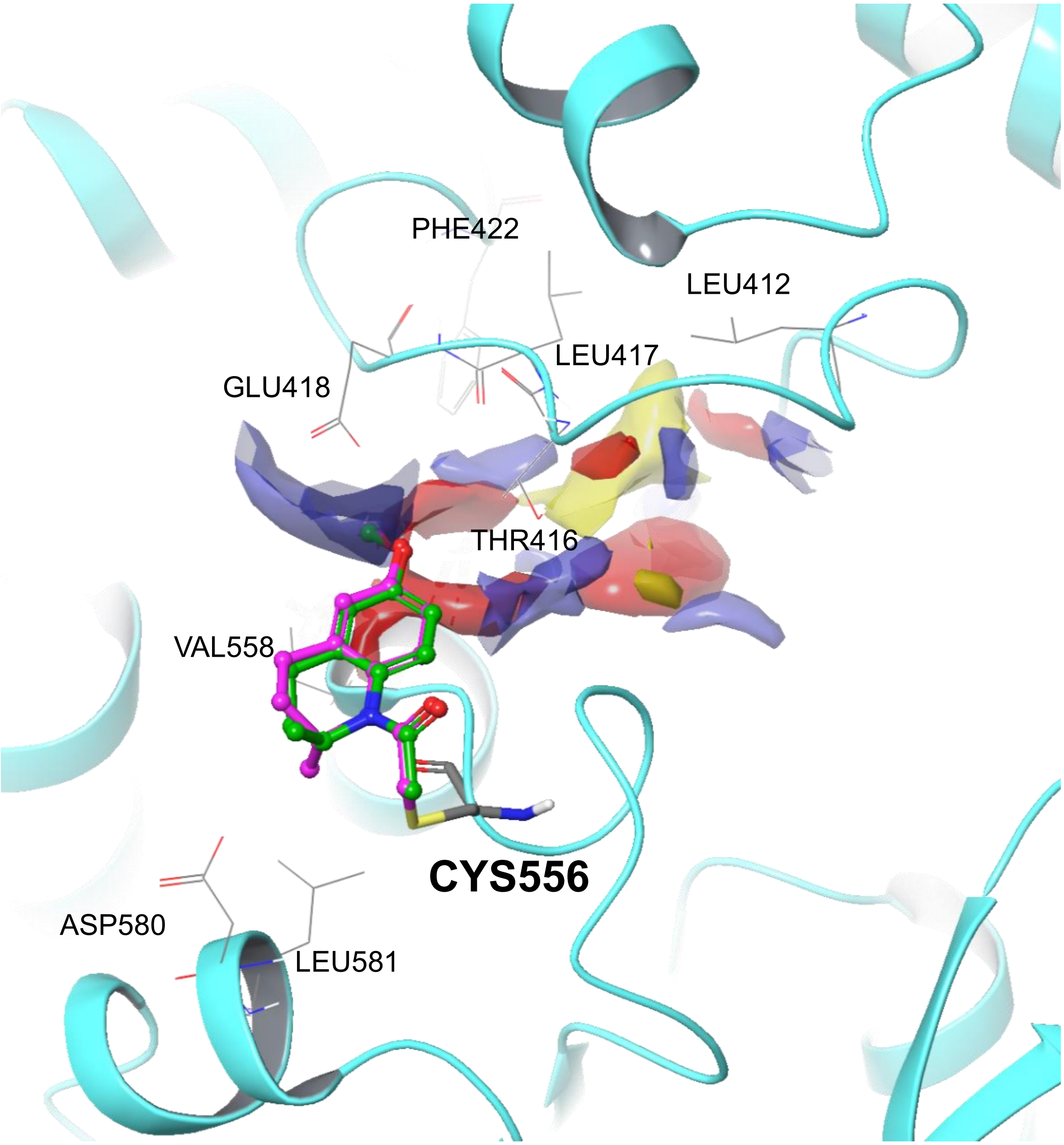


**Figure S5:** Nsp13 Covalent docking model of **3a** (purple) and **3b** (green) and binding site characterization. Yellow indicates the hydrophobic region. Red and blue indicate H-bond acceptor favorable region and H-bond donor favorable region, respectively.

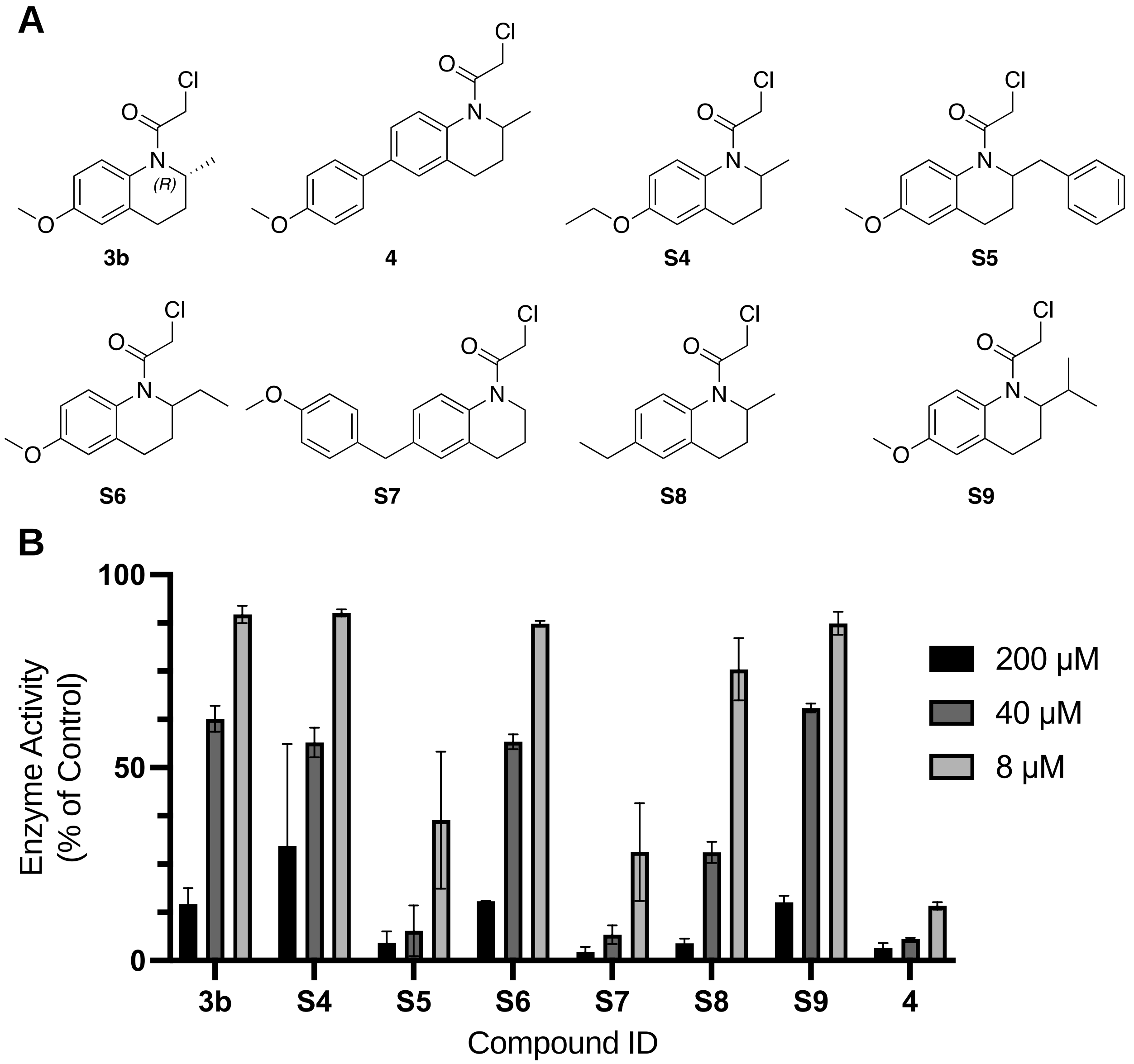


**Figure S6:** Compounds tested in second analog screen and inhibitory activity. (a) Chemical structures of compounds tested. (b) Testing of compound **3b** analogs. (8h incubation, 4$^{o}$C)

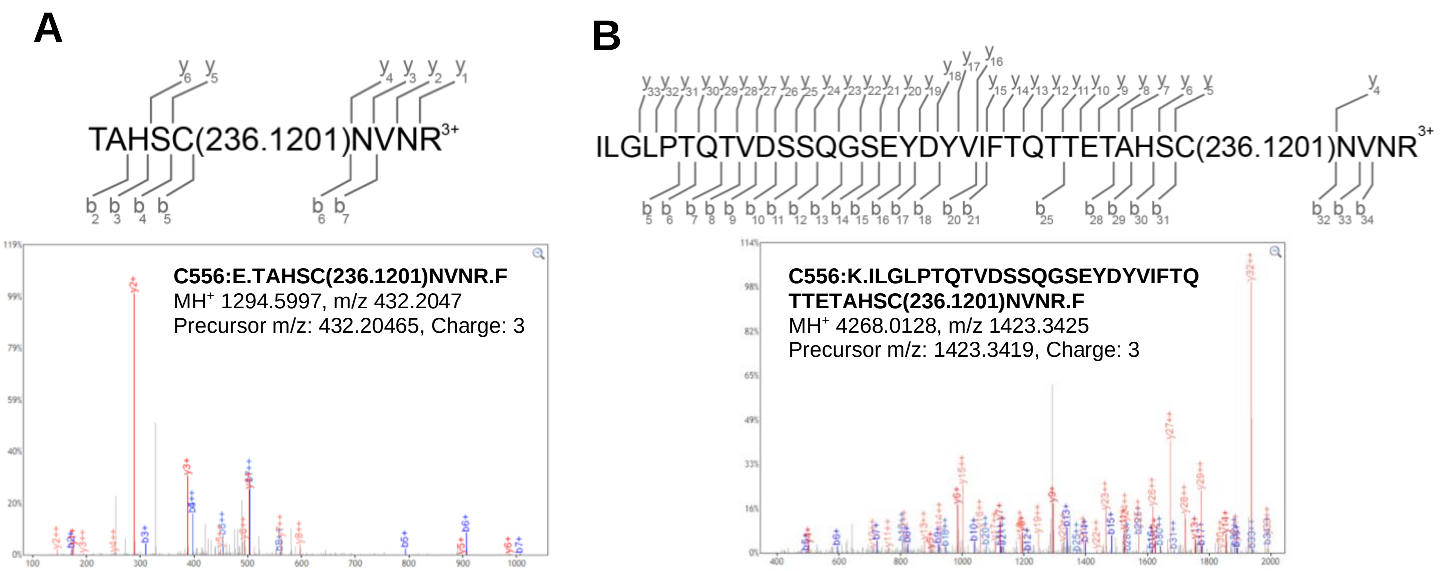


**Figure S7:** Site-mapping MS analysis of nsp13$^{wt}$ treated with compound **4b** (20 µM, 4h, 4$^{o}$C) a), b) Representative MS$^2$ spectra corresponding to peptides liganded by compound **4b** at C556. Note: multiple cleavage peptides detected based on inclusion of GluC protease during sample preparation (see methods).

| Nsp13 Cys Residue | Liganded Peptide Counts | Unliganded Peptide Counts | Total Peptide Counts | Liganding Efficiency' (%) |
|---|---|---|---|---|
| 309 | 1 | 117 | 118 | 1 |
| 342 | 1 | 22 | 23 | 4 |
| 426 | 3 | 72 | 75 | 4 |
| 444 | 3 | 179 | 182 | 2 |
| **556** | **21** | **3** | **24** | **88** |

**Table S4:** Residues of nsp13$^{wt}$ liganded by compound **4b.**

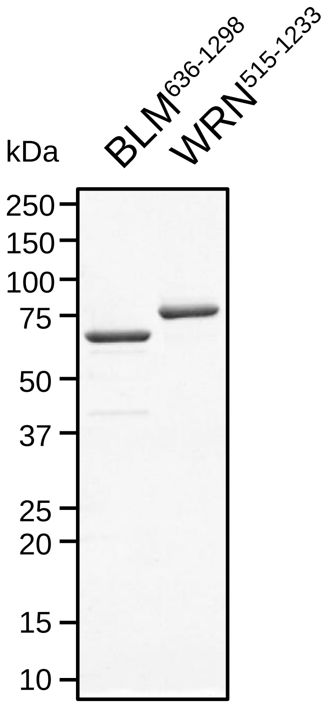


**Figure S8:** Purification of BLM$^{636\text{-}1298}$ and WRN$^{515\text{-}1233}$ constructs. SDS-PAGE gel of purified recombinant BLM and WRN.
